# 3D printed scaffold combined to 2D osteoinductive coatings to repair a critical-size mandibular bone defect

**DOI:** 10.1101/2020.12.08.415778

**Authors:** Michael Bouyer, Charlotte Garot, Paul Machillot, Julien Vollaire, Vincent Fitzpatrick, Sanela Morand, Jean Boutonnat, Véronique Josserand, Georges Bettega, Catherine Picart

**Affiliations:** CEA, CNRS, Université de Grenoble Alpes, ERL5000 BRM, IRIG Institute, 17 rue des Martyrs, F-38054, Grenoble, France; CNRS and Grenoble Institute of Engineering, UMR5628, LMGP, 3 parvis Louis Néel, F-38016 Grenoble, France; Institut Universitaire de France, 1 rue Descartes, 75231 Paris Cedex 05, France; INSERM U1209, Institut Albert Bonniot, F-38000 Grenoble, France; Université Grenoble Alpes, Institut Albert Bonniot, F-38000 Grenoble, France; Unité médico-technique d’Histologie Cytologie expérimentale, Faculté de Médecine, Université Joseph Fourier, 38700 La Tronche, France; Département d’Anatomie et Cytologie Pathologique, Institut de biologie et de pathologie, Centre Hospitalier Universitaire de Grenoble, France; Service de chirurgie maxillo-faciale, Centre Hospitalier Annecy Genevois, 1 avenue de l’hôpital –74370 Epagny Metz-Tessy, France; Clinique Générale d’Annecy, 4 chemin de la tour la reine –74000 Annecy, France

**Author notes:** Co-last and co-corresponding authors. co-first authors (Michael Bouyer, Charlotte Garot).

**Keywords:** Scaffolds, bone morphogenetic protein 2 (BMP-2), tissue engineering, 3D printing, bone regeneration, critical-size bone defects

## Abstract

the reconstruction of large bone defects (12 cm^3^) remains a challenge for clinicians. We developed a new critical-size mandibular bone defect model on a mini-pig, close to human clinical issues. We analyzed the bone reconstruction obtained by a 3D printed scaffold made of clinical-grade PLA, coated with a polyelectrolyte film delivering an osteogenic bioactive molecule (BMP-2). We compared the results (CT-scan, μCT, histology) to the gold standard solution, bone autograft. We demonstrated that the dose of BMP-2 delivered from the scaffold significantly influenced the amount of regenerated bone and the repair kinetics, with a clear BMP-2 dose-dependence. Bone was homogeneously formed inside the scaffold without ectopic bone formation. The bone repair was as good as for the bone autograft. The BMP-2 doses applied in our study were reduced 20 to 75-fold compared to the commercial collagen sponges used in the current clinical applications, without any adverse effects. 3D printed PLA scaffolds loaded with reduced doses of BMP-2 can be a safe and simple solution for large bone defects faced in the clinic.

## Introduction

To date, autologous bone graft remains the major clinical solution to treat extensive bone loss and trauma [1], but it is hampered by several drawbacks, including limited availability, pain for the patient, additional healing time and donor site morbidity. Tissue engineering using synthetic scaffolds, bioactive factors and/or stem cells offers alternative therapeutic strategies and holds promise for bone regeneration [2], but the repair of large bone defects (around 5 cm^3^) remains challenging [3]. In particular, for large bone defects, a structural synthetic scaffold may not be enough to support complete regeneration. Ceramics, notably composites of hydroxyapatite (HAP) and tricalcium phosphate (TCP) are the most biomimetic scaffolds [4], but are brittle and exhibit some variable biodegradability. Besides, they induce a basic level of bone formation [5]. Metals such as titanium are interesting for their mechanical properties [6] but can yield to stress shielding and are not biodegradable. The use of polymers has expanded, in view of their versatility, tunable mechanical properties and biodegradability. To date, polycaprolactone (PCL) and poly(lactic acid) (PLA) derivatives are the most widely used in bone tissue engineering [7].

Very interestingly, recent developments in additive manufacturing enable the design of custom-made 3D architectured scaffolds that can be adapted to the defect size [8] and are easier to implement from a regulatory perspective [9]. Polymers are particularly well-suited for additive manufacturing of scaffolds [10]. They can be manufactured in the form of filaments and be 3D printed using several techniques, including fused deposition modeling (FDM) [2, 11]. The 3D architectured scaffold plays the role of a space filler that should be mechanically stable enough to enable bone ingrowth inside the pores of the scaffold.

However, for large defect areas, a structural scaffold may not be enough to support complete regeneration. In these cases, stem cells or exogenous factors can be added to the scaffold in order to enhance regeneration [12]. Using stem cells in combination with scaffolds appears to have potential in view of their secretion of factors [13], but is more complicated to set up since different steps are required to harvest the cells from the patients, expand them in culture, and finally implant them back into the patient. As an alternative to stem cell implantation, the use of growth factors aims to recruit stem cells directly to the site of implantation. So far, bone morphogenetic protein 2 (BMP-2) has been the most widely studied clinically-approved protein in view of its ability to directly target BMP receptors at the cell surface and trigger stem cell differentiation in bone [14, 15]. BMP-7 has also been used in combination with TCP/PCL scaffolds [5]. The challenge is to optimize the dose of BMPs to avoid possible side effects [12, 16] that can lead to inflammation and ectopic bone [17]. Recently, a BMP/activin A chimera (BV-265) has been developed with increased receptor binding. [18]. Used in a composite matrix made of HAP granules and collagen I, BV-265 improved bone repair in nonhuman primate bone defect models, and allowed a 30-fold decrease in dose compared to BMP-2.

As an alternative to the incorporation of BMPs inside a carrier, surface coatings appear interesting [19, 20] in that they enable to decouple the 3D scaffold architecture from the 2D osteoinductive coatings. In our previous study [21], we showed that it is possible to repair a critical-size femoral bone defect in rats by combining a polymeric scaffold (3D hollow tube) with an osteoinductive surface coating using a polyelectrolyte film coating as a BMP-2 carrier.

An important step toward the clinical translation of a tissue-engineered construct is its pre-clinical testing on critical-size bone defects in large animals [2], which are needed to repair large bone volume defects (typically bone defects with a volume above 5 cm^3^). To date, preclinical models still need to be improved [22] and are limited when it comes to the clinical translation of results [3, 22]. Less than 13% of the studies are done in large animals (sheep, goat, dog and pig) and the vast majority do not use skeletally-mature large animals [3]. Here, we used a 3D polymeric scaffold made of clinical-grade PLA combined with an osteoinductive surface coating containing a tunable dose of BMP-2 to repair a critical-size bone defect in pig mandibles on mature animals. We designed and printed the 3D scaffold to fit a parallelepipedal volumetric bone defect of 12 cm^3^. We coated its surface with a polyelectrolyte film as a BMP-2 carrier, which enables to deliver a tunable dose of BMP-2 locally, from the surface of the scaffold. The film-coated architectured scaffolds were implanted in a full thickness mandibular bone defect for three months. They induced bone growth inside the pores of the scaffold in a homogeneous manner. Bone repair was BMP-2 dose-dependent and was as good as the autograft control. These results support further evaluations of these new osteoinductive medical devices in human bone repair clinical trials.

## Materials and Methods

### PLA scaffold preparation, polyelectrolyte multilayer film coating

Parallelepipedal 1 × 3 × 4 cm biodegradable and bioresorbable scaffolds made of medical grade poly(lactic) acid (Poly-Med, Inc, Lactoprene® 100M Monofilament 1.75 mm) and fabricated by fused deposition modeling (3DXP - One) were manufactured. PLA filaments of about 400 μm in diameter were deposited following a +45°/-45° pattern with a height of 0.2 mm and an interspacing distance of ∼2 mm. The scaffold had a porosity of 85%, with fully interconnected pores. After the fabrication and before the coating with polyelectrolye films, the scaffolds were stored away from moisture in a desiccator with a silica gel.

In order to render the scaffold osteoinductive, it was coated with a polyelectrolyte film made of 24 alternating layer pairs of PLL and HA, as previously described [21]. Briefly, the polyelectrolyte films were deposited layer-by-layer with a DR3 dip coating robot (Kirstein et Riegler GmbH). A first layer of poly(ethyleneimine) (Sigma-Alrich) was deposited using a concentration of 5 mg/mL, followed by alternated layers of hyaluronic acid (HA at 1 mg/mL, Lifecore, USA) and poly(L-lysine) (PLL at 0.5 mg/mL, Sigma, France). The film crosslinking level was controlled by incubating the coated scaffolds in 30 or 70 mg/mL 1-ethyl-3-(3-dimethylaminopropyl)carbodiimide (EDC, Sigma, France). The film crosslinking level has an effect on the film stiffness as shown in previous studies [23], EDC30 films being softer than EDC70 films. After UV-sterilization of the film-coated implants, BMP-2 (InductOs, Medtronic) was post-loaded in the polyelectrolyte films at increasing loading concentrations of 20, 50, or 110 μg/cm^3^, as previously described [21, 24]. Finally, the osteoinductive-coated scaffolds were rinsed, dried, and stored away from moisture in a desiccator with a silica gel until implantation.

### Characterization of polyelectrolyte films and quantification of BMP-2 loading

Fluorescence microscopy and scanning electronic microscopy (SEM) were used to characterize the film coating on the scaffolds. For SEM, the air-dried polyelectrolyte films coated on the PLA scaffolds were imaged) using a FEI-Quanta 250 SEM-FEG in high vacuum at 15 keV using the Everhart-Thornley detector, as previously described [21, 25]. To assess the effective coating of the film, the film was scratched with a needle and observed using SEM. Regarding fluorescence miscrocopy, the film-coated scaffolds were labeled with PLL^FITC^, as previously described [26]. They were imaged using a Leica Macrofluo (Z16 Apo) fluorescence system using a 0.8X objective [25] to assess the global homogeneity of the coating and a Zeiss LSM 700 confocal microscope with a 10X objective to assess the homogeneous coating of the film.

The quantification of BMP-2 initially loaded in the polyelectrolyte film (deposited at the bottom of a 96-well microplate) was done using a micro bicinchoninic acid assay (microBCA) test for low BMP-2 concentration (BMP20) while Nanodrop (Thermofisher) was used for higher concentrations. The concentration of BMP-2 in the loading solution was measured initially and then after incubation with the film-coated scaffold. The loaded amount, corresponding to the difference between these two values was also expressed as μg of protein per volume of scaffold (μg/cm^3^).The quantity of BMP-2 effectively loaded onto the scaffolds was then deduced from this quantification (**Table 1**). The % of BMP-2 release in vitro after several washes with a physiological buffer (HEPES-NaCl) was determined by fluorescence spectrometry using BMP-2 carboxyfluorescein (BMP-2^CF^).

**Table 1.** Quantification of the effective amount of BMP-2 loaded in the film-coated architectured scaffold. The total volume of the scaffold was 12 cm^3^ and its effective surface estimated from the design was 144 cm^2^. We targeted a BMP-2 loading (unit mass per unit of scaffold volume in μg/cm^3^). The total amount of BMP-2 effectively loaded was calculated for each implant and reported in “mass per volume of implant” (μg/cm^3^)

|  | Targeted BMP-2 loading (µg/cm<sup>3</sup>) | Total BMP-2 Loaded (µg/implant) | Volumetric dose (µg/cm<sup>3</sup>) |
| --- | --- | --- | --- |
| BMP20 (n=2) | 20 | 240 | 20 |
| BMP110 (n=2) | 110 | 870 | 72.5 |

Main experiment
|  | Targeted BMP-2 loading (µg/cm<sup>3</sup>) | Total BMP-2 Loaded (µg/implant) | Volumetric dose (µg/cm<sup>3</sup>) |
| --- | --- | --- | --- |
| BMP50 (n=6) | 50 | 326 ± 80 | 27 |
| BMP110 (n=5) | 110 | 1000 ± 64 | 83 |

### In vivo critical-sized mandibular defect in minipigs

Mature female minipigs (lineage FBM “down-sized”, age over 24 months) weighing 43 to 69.5 kg (mean 56.3 kg), were included in the study, which was approved by the animal ethics committee (N° *APAFIS#2876-2015111616474259 v2*). They were acclimated for 2 weeks before surgery and immediately accustomed to a smooth diet. Animals were given antibiotics 2 days before surgery (Amoxicillin and Clavulanic acid at 12.5 mg/kg). They were fasting almost six hours before surgery. They were given pre-anesthetic medication in the form of Atropin 0.04mg/kg, Azaperone 2 mg/kg by intramuscular injection and Morphine 0.2 mg/kg by subcutaneous injection. Induction was performed with Tiletamine and Zolazepam (3 to 5 mg/kg). After oral intubation, anesthesia was maintained with isoflurane. Animals were placed in supine position, the mandibular area was shaved and prepared with an iodine scrub. The mandibular body was exposed via a submandibular approach, leaving the periosteum on the bone. Bone resection was made with an oscillating saw, creating a full thickness bone defect of 4 x 3 cm, including the periosteum, and penetrating the mandibular nerve canal. This resection was standardized using a phantom (an uncoated implant) to delimit the perimeter of resection. Two titanium reconstructive plates (Stryker 2.8 system, Freiburg, Germany) and twelve 2.7 mm diameter screws were used to stabilize the mandible by triangulation, before insertion of the implant in the defect (Stryker Leibinger GmbH & Co. Freiburg, Germany).

The implant was fixed on the titanium plate using 2/0 nylon stitches (**Figure 1**). When a bone graft was used (positive control group), it was harvested from the iliac bone and fixed on the plate using two screws. The wound was closed in three layers without drainage. Both sides of the mandible were operated on in the same way.

**FIGURE 1.**
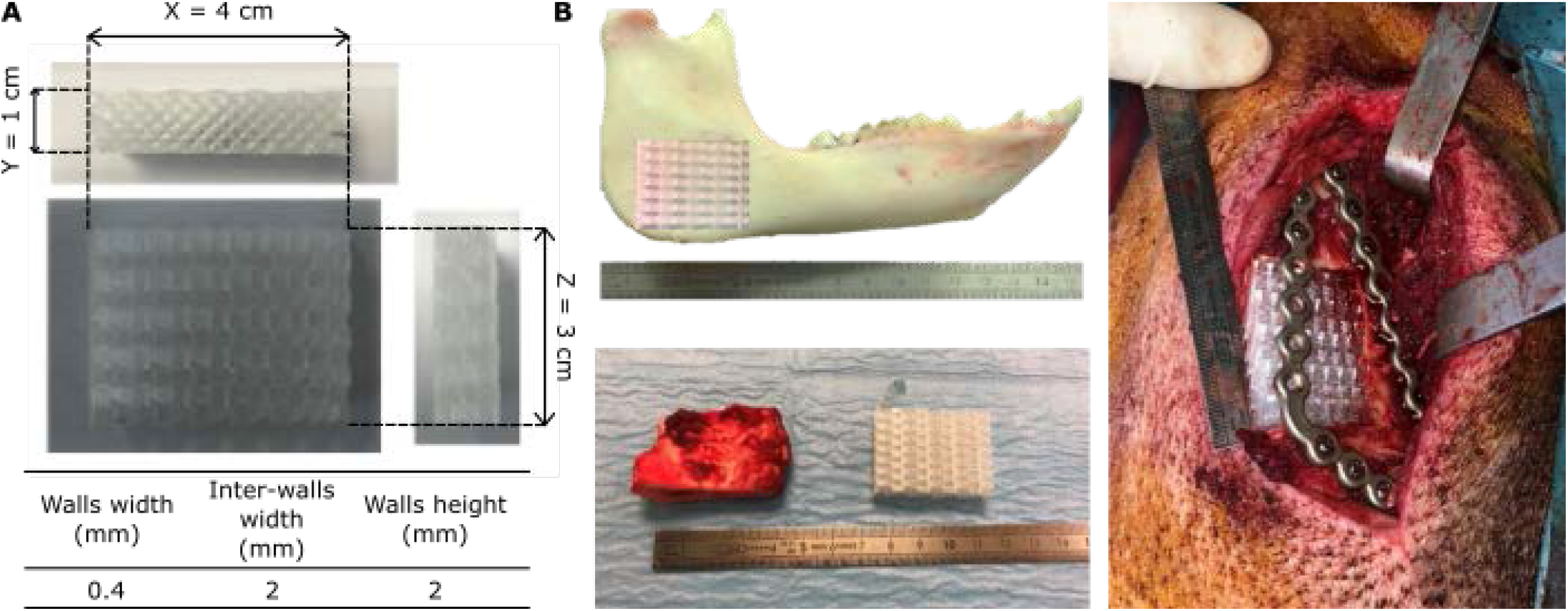
3D printed PLA scaffold and experimental design of the in vivo experiments in the minipig mandible. (A) Parallelepipedal scaffolds (4 x 3 x 1 cm, total volume of 12 cm^3^) were custom-made with medical-grade PLA using a fused deposition melting 3D printer. PLA filaments of ∼ 400 μm in diameter were deposited in a +45°/-45° pattern, with an interspacing of 2 mm, resulting in a scaffold with a porosity of 85% with fully interconnected pores. This pore size was designed to promote cell adhesion and the transport of nutrients into the implant. (B) Implantation in minipig mandibles. First, a bone defect was created into the minipig mandible and the bone was cut. Two 2.8 mm plates were fixed with almost twelve screws of length 2.7 mm and the implant was positioned in between with nylon stiches.

Aftercare included Morphine 0.2 mg/kg injected subcutaneously according to clinical symptoms, a fentanyl patch (50 μg/h) changed every 3 days for a period depending on the residual pain, antibiotics (Amoxicillin and Clavulanic acid 12.5 mg/kg) intra-orally for 15 days, meloxicam 0.4 mg/kg for 3 days. The animals were followed for 13 weeks in individual boxes. A veterinarian clinically evaluated the animals every day, three times per day during the first 2 days after surgery and once per day after. Analgesia was adapted according to symptoms. The animals were weighed once a week. Special attention was given to re-feeding and weight evolution. Blood analyses were made before surgery and every week until euthanasia to test liver, kidney, hematological functions and inflammation. Complete blood count, haptoglobin and protein electrophoresis were measured to assess inflammation and hemostasis, while aspartate aminotransferases (ASAT), alanine aminotransferases (ALAT), alkaline phosphatase (ALP), gamma-glutamine transferase (CGT) and bilirubinemia were used to assess hepatic function. Serum creatinine and urea were quantified to assess renal function.

First, in a preliminary experiment on 3 minipigs, we screened 6 conditions (*n*=1): two crosslinking levels EDC30 and EDC70, two BMP-2 initial loading concentrations of 20 and 110 μg/cm^3^, and two negative controls: i) empty defect to assess the critical size of the defect, and ii) film-coated scaffold (EDC70) without BMP-2. Each implant was randomized, implanted, and analyzed in a blind manner using CT-scans, the polyelectrolyte films being macroscopically indistinguishable. All analyses of experimental groups were also made in a blind manner.

In a second main experiment, two conditions were further studied in larger groups with an intermediate dose of BMP-2 (loading at 50 μg/ cm^3^ and 110 μg/ cm^3^) (*n*=6 for low BMP-2 dose, n=5 for high dose) to perform statistical analysis, another negative control was added (EDC30 film without BMP-2), and bone autograft (n=4) was added as positive control (see **Table 2** for all experimental conditions).

**Table 2.** Experimental conditions and total number of minipigs studied per group in the preliminary and second main experiment. Three BMP-2 doses (in µg/cm^3^ of scaffold) and two film crosslinking level were studied (EDC30 and EDC70) in two successive experiments: a preliminary one to study two BMP doses (BMP20 and BMP110) and compare EDC30 and EDC70 films, and test two controls no PLA scaffold to confirm the critical size of the defect, and film-coated PLA (EDC70) without BMP-2 (n=1 for all). In the second main experiment, the conditions were selected based on the results of the preliminary experiments: EDC30 films were studied for two BMP-2 doses (BMP50, *n*=6 and BMP100, *n*=5). A positive control group was added (bone autograft, *n*=4) and a negative control (EDC30 film coated PLA-implant). In this table, the first number on the left of the “+” refers to the number of sample in the preliminary experiment while the second number, refers to the number of sample in the second main experiment.

| Film condition | BMP-2 targeted amount<br>(in $\mu\text{g}$ of BMP-2 / $\text{cm}^3$ of scaffold) | | | |
| --- | --- | --- | --- | --- |
|  | BMP0 | BMP20 | BMP50 | BMP110 |
| No implant<br>(negative control) | 1+0 | - | - | - |
| EDC30 | 0+1 | 1+0 | 0+6 | 1+5 |
| EDC70 | 1+0 | 1+0 | 0+0 | 1+0 |
| Bone Autograft<br>(BG, positive control) | 0+4 | - | - | - |

After the last CT-scan (D91), the animals were euthanized by Pentobarbital injection. The skin around the mandible was examined attentively to notice eventual complications (fistulas, collection, inflammation…). Using the same submandibular approach, the titanium plates and screws were removed. On each side, a full thickness sample of bone was removed using a surgical guide (**Figure 1**), to get a margin of 1 cm native bone all around the initial implant. The bone pieces were fixed in 4% neutral buffered formalin (Sigma, France) for one week at 4°C.

### CT-scan and micro-CT analysis

Assessment of bone formation was made by CT-scan after ∼15, 30, 50 and 90 days. These radiographies were acquired using a helicoidal BrightSpeed 16 scanner (General Electric) under gaseous anesthesia.

First, a CT-scan score was defined by the authors as considering four criteria: the percentage of filling of the porous implant (F), the homogeneity of the newly formed bone (H), the ability to distinguish between cortical and cancellous bones (D) and the amount of ectopic bone (E). Each criterion was evaluated in a blind manner by four clinicians using a score between 0 and 4 (0 being the lowest grade and 4 the highest). Then, the global score (S) was calculated by each clinician and for each CT-scan made during the follow-up period:

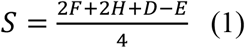

the criterion F and H related to bone growth being rated with a dooble weight. Finally, the global score was then represented as the mean score ± SD of the four scores given by each clinician independently.

Second, quantitative analysis of bone growth was also performed using the software InVesalius 3 (Centro de Tecnologia da Informação Renato Archer, CTI). The volume of interest (VOI) was defined as the total volume of newly formed bone inside and around the bone defect. Bone autograft was used as control. Based on the visual observations, we decided to segment and define two types of bones: poorly and highly mineralized. To quantify these two fractions, the VOI was segmented using a global threshold, namely between 230 and 629 HU for poorly mineralized bone and values > 630 HU for highly mineralized bone. The volumes of the newly formed bone were expressed in mm^3^. For the quantification of the bone volume outside the implant (named “ectopic bone”), the VOI was adjusted to the volume of the scaffold. Then, the volume of bone inside this new VOI was subtracted from the total bone volume formed.

Third, micro-computed tomography (μCT) imaging was performed on fixed mandibular explants after 90 days, using high resolution scanner (vivaCT 40, ScancoMedical, Switzerland), as previously described [21]. The acquisition parameters were set at 70 kV with an intensity of 114 μA, a 100-ms integration time and an isotropic voxel size of 76 μm. The volume of interest (VOI) was defined as the volume of the initial bone defect in the mandible (3 cm x 4 cm x 1 cm). Bone volume was determined after segmentation using threshold values (438-2730 mg HA/cm^3^) and Gaussian filter (sigma 0.8, support 1). Bone Mineral Density (BMD, mg HA/cm^3^) and the ratio of bone volume divided by total VOI (BV/TV) were calculated.

### Bone homogeneity score

The implant of well-defined dimensions (3 cm x 4 cm x 1 cm, total volume TV of 12 cm^3^) was taken as the region of interest (ROI). It was separated into ten slices of equal thickness along each axis (X, Y, Z). For each slice, the bone volume ratio (BVr=BV/TV) was calculated as the bone volume into one slice (BVs) divided by the volume of the slice of interest (corresponding to TV/10). This quantification was done for each axis (**Figure SI 9**): the standard deviation (SD) of BVr was calculated and the homogeneity score was defined as the sum of the three SDs over X, Y and Z axes.

### Histology and histomorphometry

The specimens were dehydrated in a graded series of alcohol and embedded in methylmethacrylate (MMA). 3 slices in the XZ plane were cut with a laser microtome (TissueSurgeon, LLS ROWIAK GmbH, Hannover, Germany) [27] and stained with Sanderson’s rapid stain and Van Gieson’s staining. Slice thickness was 10 to 100 μm. Sections were imaged using a slide scanner. Histological examination was performed by a pathologist using a Leica microscope using transmitted or polarized light. The histomorphometry analysis was performed in a blind manner by three independent operators. The operators gave a visual approximation of the bone area to total area ratio since a systemic analysis was not possible due to the high differences in staining between the different sections, named here S1, S2, and S3.

### Statistical analysis

OriginPro (OriginLab), Excel (Microsoft Office) and R for mac OS X (R foundation for statistical computing, CRAN) were used for all analyses. Data were expressed as mean ± standard deviation. Non-parametric data were presented by median and interquartile range. Differences between groups were assessed by analysis of variance (ANOVA) and Bonferroni post-hoc analysis or Student’s t-test. Differences between groups at p-values < 0.05 (*) and p < 0.01 (**) were considered as significant.

## Results

### 3D architectured scaffolds combined with an osteoinductive surface coating to repair a critical size-mandibular bone defect

Parallelepipedal scaffolds (4 x 3 x 1 cm, for a total volume of 12 cm^3^) were made of clinical-grade PLA and custom-fabricated using fused deposition modeling (FDM). The PLA filaments of ∼ 400 μm in diameter were deposited following a +45°/-45° pattern with an interspacing of 2 mm, resulting in a scaffold porosity of 85% with fully interconnected pores (**Figure 1A**) to enable the transport of fluids and nutrients in the core of the scaffold. This choice has been made after testing in laboratory several types of scaffold architectures (data not shown). The variations concerned their trabecular thickness from 1 to 2.5 mm, porosity from low to high porosity, and geometry. We qualitatively tested their compression resistance, surface state and fixation possibilities (suturing). We also reasoned based on our preliminary study using film-coated PLGA hollow cylinders that showed that 3 mm diameter cylinders enabled bone formation inside the empty space inside the cylinder [21].

As an implantation site, we chose to create a large critical-size pig mandibular defect (12 cm^3^) (**Figure 1B**) due to the fact that the pig mandible mimics the human mandible [28] [29]. This was a full thickness defect on the basilar border of the mandibular angle, scarifying the alveolar bundle. We chose a posterior location since it has been shown that the extent of defect regeneration from spontaneous healing was significantly less in the posterior than in the anterior mandibular defects [30]. Buccal and vestibular periosteum were removed around the defect to avoid spontaneous ossification. Additionally, the animals were mature, older than 24 months, without any remaining growth potential. We selected this age as our previous preliminary study on four minipigs aged six to eight months has shown some spontaneous ossification of the defect spreading from the edges (data not shown). Thus, choosing animals older than 24 months ensured that bone maturation had occurred and that new bone formation would not result from the endogenous bone formation resulting from the animal growth. The implant was inserted in the defect and fixed using nylon stitches on the two 2.8 mm thick titanium plates used to stabilize the mandible by triangulation; twelve screws fixed the plates (**Figure 1B**).

The homogeneous coating of the scaffold by the polyelectrolyte film was visualized by fluorescent labeling of the film with PLL^FITC^ and imaging using a fluorescence macroscope (**Figure 2A**) and confocal microscopy (**Figure 2B**). As observed by scanning electron microscopy after scratching the film using a needle, the polyelectrolyte film fully coated the PLA filaments (**Figure 2C**). The amount of BMP-2 initially loaded in the polyelectrolyte film and the % of release in vitro were determined by fluorescence spectrometry using fluorescently-labelled BMP-2 (**Figure 2D and Figure 2E**). This in vitro release study allows to have a general idea of the initial release, though the in vivo behavior may be different on the long term. The amount of BMP-2 increases with the initial concentration of BMP-2 in the loading solution, but it reaches a plateau faster for the EDC70 film (**Figure 2D**). The amount of BMP-2 released from the film is higher for the EDC30 film: the maximum percentage of BMP-2 released from the film is ∼50% for EDC30 films, compared to ∼20% for EDC70 films (**Figure 2E**).

**FIGURE 2.**
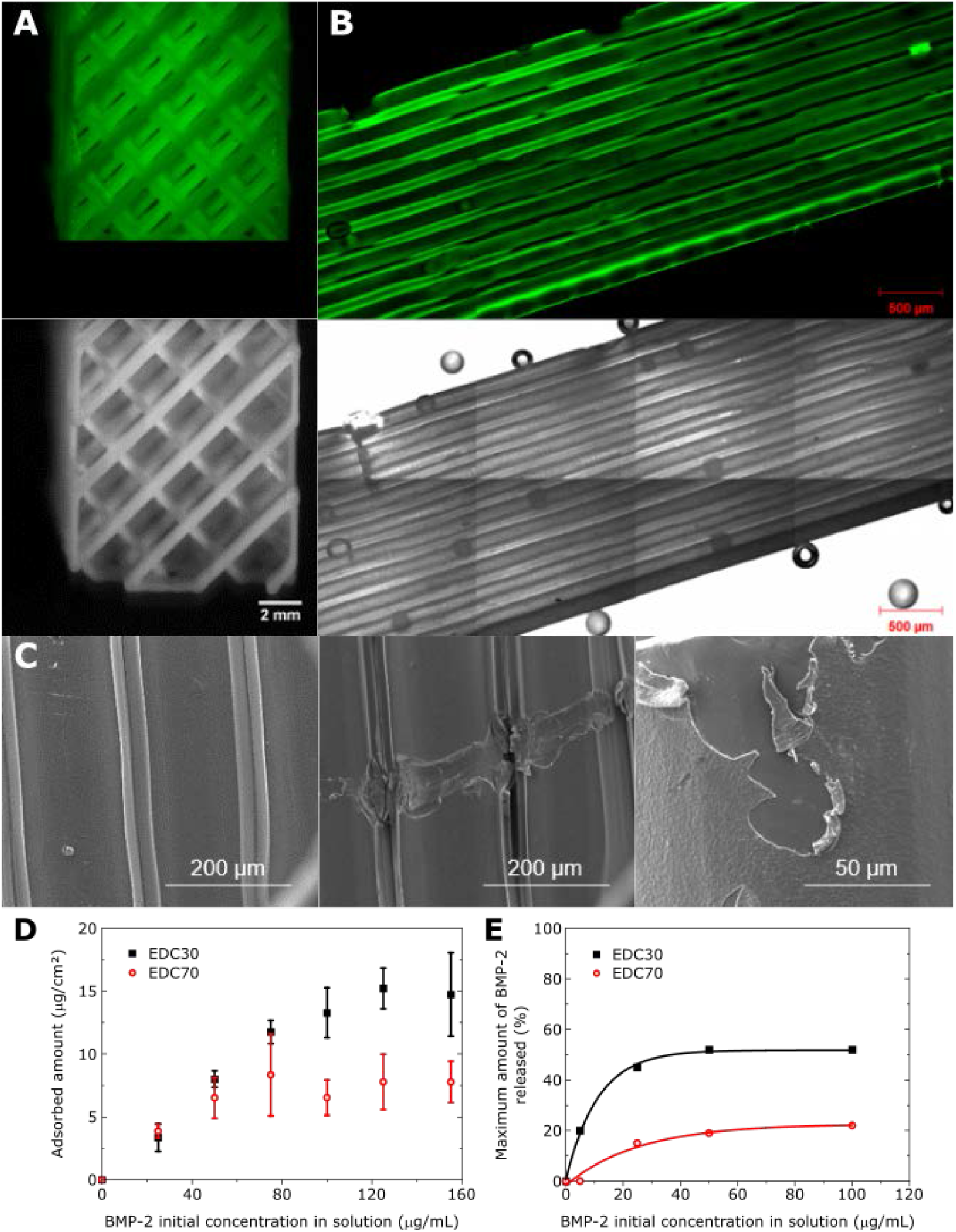
Imaging of the film-coated 3D printed PLA scaffolds. (A) Macroscope imaging of a (PLL/HA)_24_ film-coated implant using fluorescence and phase contrast imaging modes. (B) Confocal imaging of a section to visualize the film-coated struts of the scaffold. (C) SEM imaging of the struts and of a scratch made using a needle, to assess the presence of the film. (D) Quantification using μBCA of BMP-2 loading in (PLL/HA)_24_ films as a function of the initial BMP-2 concentration in the loading solution for two crosslinking levels EDC30 and EDC70. (E) Quantification of the BMP-2 release (expressed in %) as a function of the initially-loaded BMP-2 concentration, for two crosslinking levels EDC30 and EDC70.

### Preliminary experiment to validate the mandibular defect model and select the film coating conditions

To optimize bone regeneration and the operating techniques, we performed a preliminary experiment in the minipig mandibles. We initially screened 4 different conditions (*n*=1 for each condition), corresponding to two film crosslinking levels (EDC30 and EDC70) and two concentrations of BMP-2 (BMP20 and BMP110) (**Table 2**). These concentrations, expressed in μg of BMP-2 per cm^3^ of scaffold, were the targeted “volumetric” concentrations of BMP-2. Knowing the effective surface of the scaffold (144 cm^2^) and the amount of BMP-2 loaded in the polyelectrolyte film (**Figure 2D**), we defined the concentration of BMP-2 in the loading solution (in μg/mL) into which the scaffold was dipped. The amounts of BMP-2 that were effectively loaded in the film-coated 3D scaffolds were quantified (**Table 1**). Two negative controls were added: an empty defect without any implant, and a defect with the film-coated implant (crosslinking level EDC70) but without BMP-2.

The animals were in good health. There was no postoperative infection, implant failure or sign of blood disorder. All the surgical procedures were uneventful and there were no surgical complications. For one implant, it was necessary to recut the anterior border of the defect in order to improve the fit of the implant. All the titanium plates were stable and fixed to the native bone (no loosening). During explantation, it was impossible to macroscopically identify the implant from bone reconstruction or from scar tissue. A complete blood analysis was performed before surgery, immediately after surgery and once a week until euthanasia (**Figure SI 1**). The objective was to assess an eventual general inflammatory reaction, or liver or kidney complications due to BMP or due to the resorption of the film and scaffold. The blood sample analysis on three different mini-pigs did not reveal any abnormality. We concluded that the scaffold with or without the film and/or BMP-2 did not cause a general inflammation, a hepatic reaction, or a renal function impairment. Computed tomography (CT) was acquired during the follow-up period (**Figure SI 2**). For each acquisition, a score of the CT-scan was given in a blind manner by four clinicians (**Figure 3A and B**). All BMP groups exhibited bone regeneration, regardless of the crosslinking extent of the film and the BMP-2 loading concentration, while the negative controls did not show bone formation (**Figure 3B and Figure SI 2**). The CT-scan score was used to calculate a plateau value (B_max_) and a characteristic time to reach the plateau (τ), by fitting an exponential function to the experimental data [21]. For the EDC30 films (**Figure 3A**), the scores steadily increased before reaching B_max_, which was higher for the BMP110 than for BMP20 (4.8 ± 0.4 versus 1.4 ± 0.0, respectively). τ was approximately 2.6 times faster for the low dose than for the high dose (21 ± 2 days versus 55 ± 10 days). In contrast, for the EDC70 films, the exponential fit to the data was poor for the highest BMP-2 concentration, and there was no clear dose-dependent trend (**Figure 3B**). For the low BMP-2 dose, B_max_ was at 3.3 ± 0.4 and τ was 25 ± 10 days.

**FIGURE 3.**
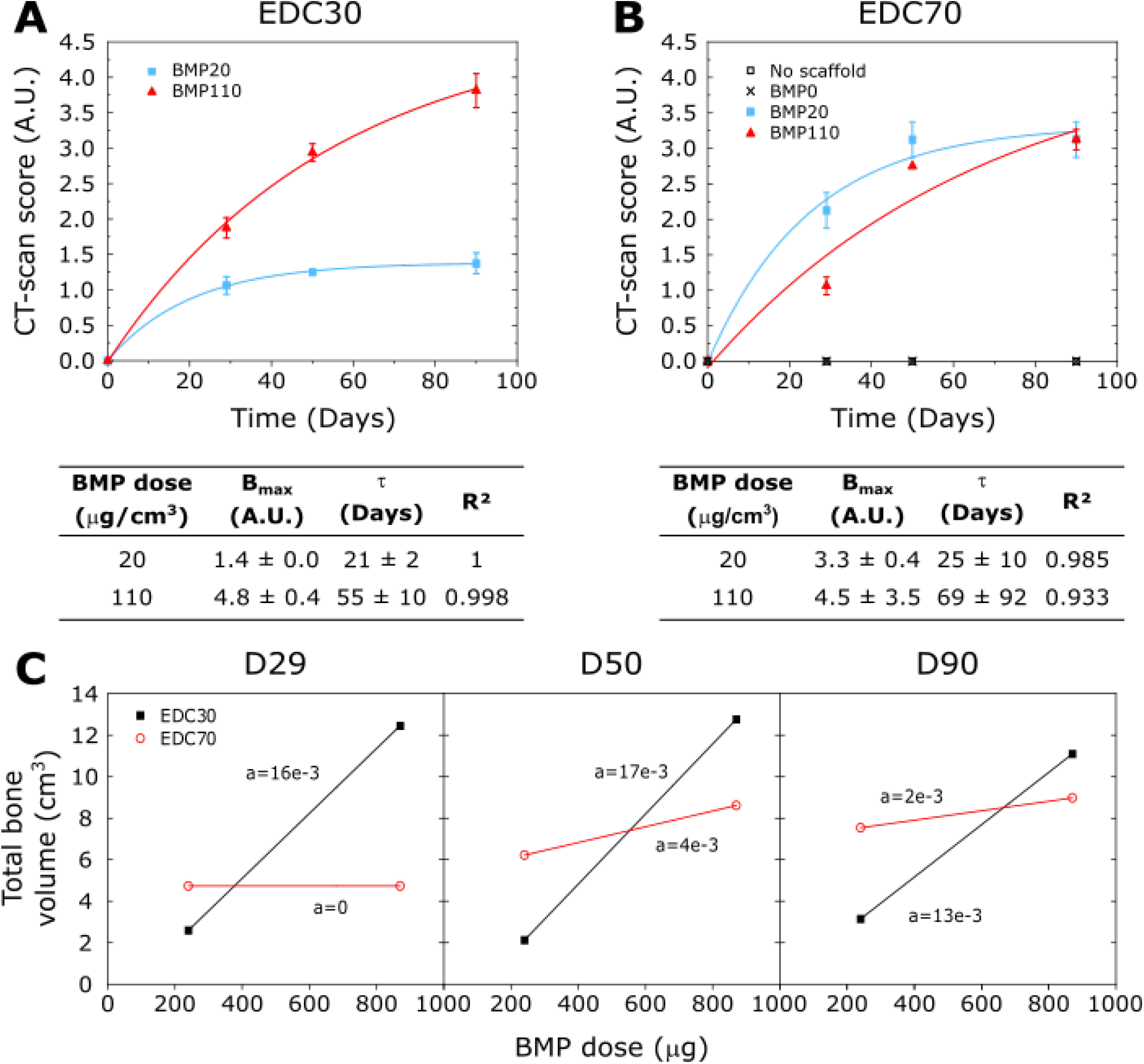
Kinetic quantification of bone formation using CT-scans obtained for the preliminary experiment with two crosslinking levels and two BMP-2 doses. EDC30 and EDC70 films loaded with BMP-2 doses (20 and 110 μg/cm^3^ of scaffold corresponding to a total dose of 240 and 870 μg per implant respectively) were compared. Two negative controls were added: empty defect (no scaffold) and film-coated implant without BMP-2. (A, B) CT-scan scores (mean of the global scores (S) given by the four clinicians ± SD) calculated from the CT scans as a function of time, and corresponding exponential fits to the data (red and blue lines) for the (A) EDC30 and (B) EDC70 groups. The plateau value (B_max_), characteristic time (τ) deduced from the fits, and fit quality R^2^ are given in the corresponding tables. (C) Total bone volume as a function of time (D29, D50 and D90) as a function of the BMP-2 dose expressed in total dose per implant (in μg), for the two crosslinked films EDC30 and EDC70. Lines have been added to guide the eye.

The quantification of the total regenerated bone volumes from the CT images for the different time points (D29, D50, and D90) confirmed the BMP-2 dose-dependence of bone repair for the EDC30 films and not for the EDC70 films (**Figure 3C**). The amount of poorly mineralized bone and highly mineralized bone were also plotted (**Figure SI 3**). The slopes of the linear fits were higher for the highly mineralized part, suggesting that this type of bone has more influence on the total bone volume than the poorly mineralized one.

μCT scans were acquired after explantation of the scaffolds. They were used to calculate the total bone volume (BV) and the bone mineral density (BMD) for all samples (**Figure SI 4)**. After 3 months, BV also exhibited a clear BMP-2 dose-dependence, independently of the EDC crosslinking level, while the BMD was rather stable. The μCT imaging of the EDC70 films confirmed that there was no visible BMP-2 dose-dependence for this film condition (**Figure SI 5**).

The associated histological examination performed by a pathologist showed no sign of foreign body reaction (**Figure SI 6**). For the low BMP-2 dose loaded on an EDC30 film-coated scaffold, very few new bone was formed and mesenchymal tissue was largely observed (**Figure SI 6A**). In contrast, the condition EDC30/BMP110 showed high bone formation with Haversian canals and blood vessels already formed (**Figure SI 6B**). The film-coated implant without BMP-2 showed no bone formation and only mesenchymal tissue could be observed (**Figure SI 6C)**. For the EDC70 film-coated implants, and independently of the BMP-2 dose, the histological sections showed bone tissue in formation (some mesenchymal tissue remained), with blood vessels and some Haversian canals already formed (**Figure SI 6D and 6E**), proving that the bone was becoming more and more mature as it was growing. The quantitative histomorphometric analysis of bone area over total area (BA/TA) confirmed a clear BMP-2 dose-dependence of bone repair for the EDC30 films with a BA/TA ratio of 1.5 ± 1% for BMP20 and 40.8 ± 1.4% for BMP110 (**Figure SI 6F**). Surprisingly, BA/TA significantly decreased with the BMP-2 dose for the EDC70 films (25.8 ± 3.8% for BMP20 versus 12.5 ± 1.5% for BMP110).

Altogether, our results established the critical size of the mandibular bone defect, the difference in bone repair kinetics depending on the BMP-2 dose, and the influence of crosslinking level of the film on the amount of newly formed bone. A clear BMP-2 dose-dependence was evidenced for the EDC30 films.

### Main experiment shows that BMP-2 influences the bone repair kinetics and the amount of highly mineralized bone

In view of the results of the preliminary experiments showing that BMP20 lead to only few bone formation, we decided to puruse the next study with only EDC30 films and selected two doses. We next repeated the experiments with more mini-pigs per condition (**Table 2**) to assess quantitatively the effect of BMP-2 dose. We decided to keep the high dose of BMP110 (n=5) and to increase the low dose of BMP-2 to BMP50 (n =6). We added the bone autograft (BG) as a positive control (*n*=4), and a film-coated scaffold without BMP-2 as a negative control (EDC30 film). An additional earlier time point (D16) was also added for the CT scan acquisitions.

Once again, there was no surgical complication. In two cases, the bone autograft was in two pieces because of the small size of the iliac bone, but in all cases the defect was completely filled. In two cases (bone autograft), a small serous collection was found around the implant, and in one case (high dose of BMP-2) a small suppurated collection with a cutaneous fistula appeared at D90.

**Figure 4** shows CT scans over time for the bone autograft (BG), the synthetic bone implant obtained for the two BMP-2 doses (BMP50 and BMP110), and the negative control (Ctrl -). At first sight, we observed that the bone autograft mineralizes over time but its total amount does not change. In contrast, the film-coated scaffold induced bone repair over time, and mineralization was also visible. The total bone volume, the poorly mineralized and highly mineralized bone volumes were quantified from the CT scans (**Figure 5A, B, C**). The total bone volume increased over time to reach a plateau, this increase being significantly higher for the high BMP-2 dose condition than for the low dose (**Figure 5A**). The bone graft volume remained constant. The amount of poorly mineralized bone quickly increased with time for both BMP50 and BMP110, but with no statistical difference (**Figure 5B**). In contrast, the amount of highly mineralized bone increased as a function of time, reaching a peak at D51 (**Figure 5C**). It was significantly higher for BMP110 than for BMP50.

**FIGURE 4.**
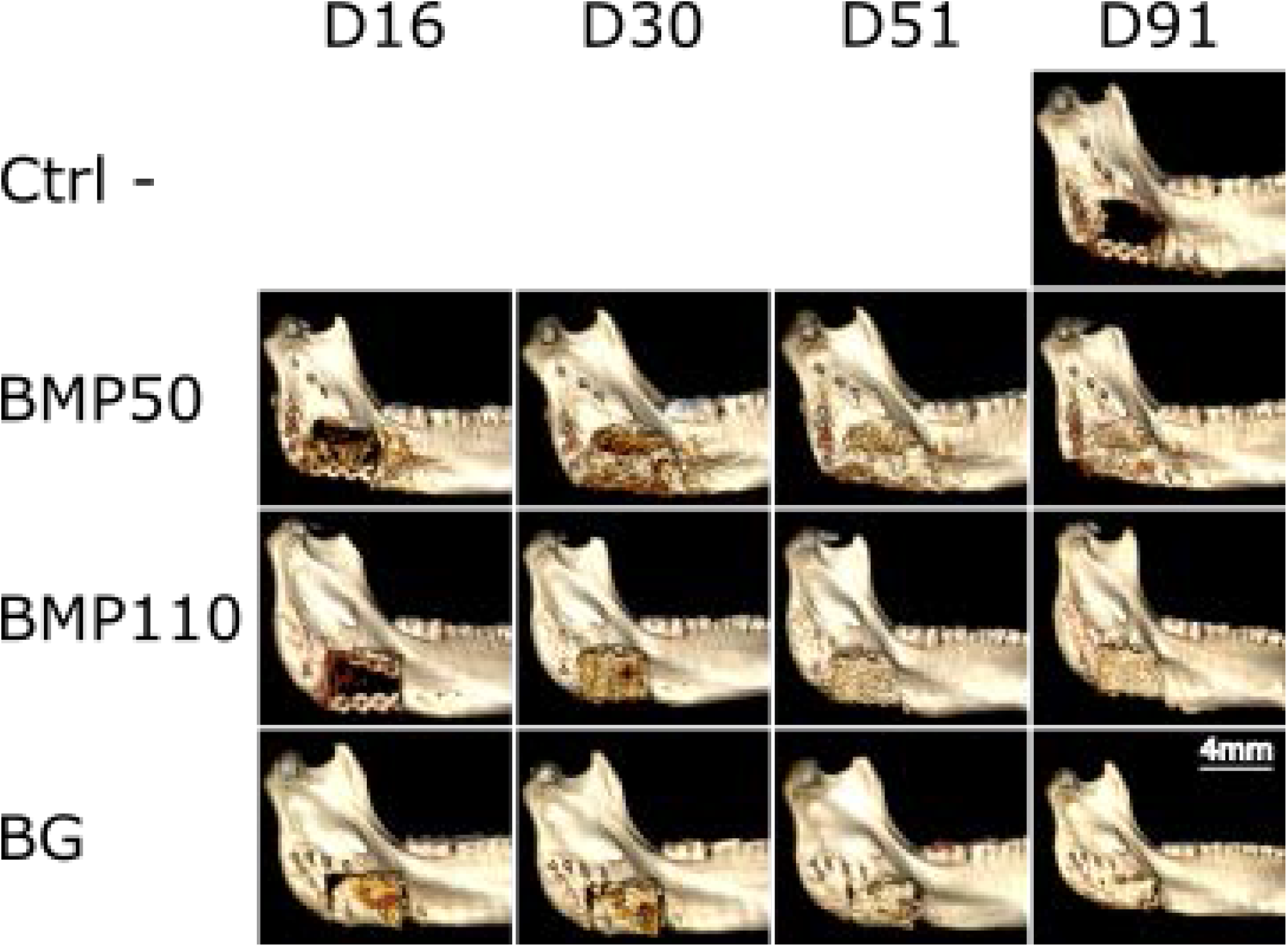
Representative 3D reconstructions of the CT scans showing the kinetics of bone regeneration for four representative conditions: negative control (Ctrl-, film-coated scaffold without BMP-2 in the film), film-coated scaffold with a low BMP-2 dose (BMP50) and a high BMP-2 dose (BMP110), and bone autograft (BG). Scale bar is 4 mm.

**FIGURE 5.**
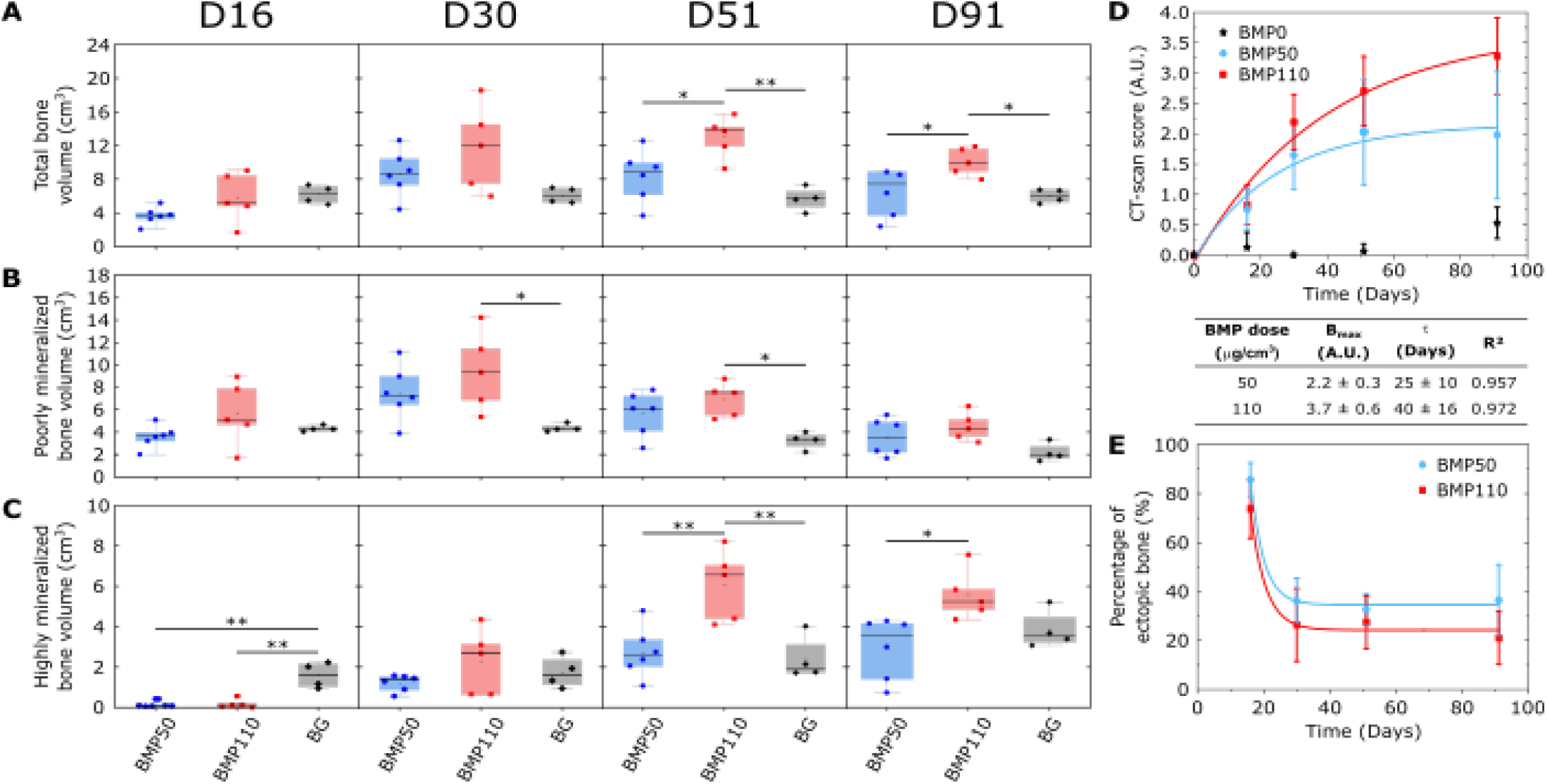
Quantitative analysis of the kinetics of bone formation followed by CT-scan for EDC30 films loaded with two doses of BMP-2. The film-coated scaffolds were loaded with BMP-2 at 50 (*n*=6) and 110 μg/cm^3^ (*n*=5) and their bone regenerative capacity was compared to bone autograft BG, (*n*=4). (A-C) Box plot representations of the total bone volume (A), poorly mineralized (B) and highly mineralized bone volumes (C) as a function of the BMP-2 dose BMP50 versus BMP110 in comparison to BG. (D) CT-scan scores as a function of time and corresponding exponential fits to the data (colored lines) for EDC30 films; corresponding plateau value (B_max_), characteristic time (τ) deduced from the fits, and fit quality R are given in the table. (E) % of bone outside the implant (named “ectopic bone”) as a function of time for BMP50 and BMP110. * p < 0.05; ** p < 0.01.

The CT-scan scores were plotted as a function of time, and the data was fitted with an exponential fit (**Figure 5D**). The score increased for the two BMP-2 doses, but again with different characteristics: B_max_ was lower for the scaffolds with BMP50 than that of the BMP110 (2.2 ± 0.3 versus 3.7 ± 0.6). τ was 1.6-fold lower for BMP50 (25 ± 10 days) than for BMP110 (40 ± 16 days).

We further analyzed the part of the regenerated bone that was forming outside of the implant (**Figure 5E**) that we call here “ectopic bone”. Initially high, the fraction of bone growing outside of implant quickly reached a plateau value at around 28 to 35%, independently of the BMP-2 dose. We noted that the dispersion of the values was slightly higher for the high dose, and also slightly higher at the day 91 endpoint.

The μCT analysis performed at day 90 after euthanasia of the minipig was used to further analyze the newly formed bone (**Figure 6 and 7**). μCT images are shown in the different planes ((X,Z), (X,Y), and (Y,Z)) for scaffolds containing increasing doses of BMP-2, taken from the experimental groups. The negative control confirms the critical size of the mandibular bone defect, and the bone autograft (BG) provides a positive reference value. For the lowest BMP-2 dose, bone formation was scarce. The amount of bone progressively increased, and the newly formed bone entirely filled the pores of the 3D scaffold in a homogeneous manner. There was no sign of excessive ectopic bone formation, even at the highest doses. This was also visible on the 3D reconstructed μCT images (**Figure 7** and **Figure SI 7**). The mean bone volume was higher for BMP110 (7.6 cm^3^) than for BMP50 (4.8 cm^3^) **(Figure 7B**), and was also higher than for the bone autograft reference. In addition, bone regeneration was more dispersed with BMP50, with a variation coefficient of 34% compared to 13% for BMP110. When plotting all the experimental bone volumes as a function of the BMP-2 total dose per implant, a linear correlation was found (**Figure 7C**). The BMD was not significantly different for the different doses but was significantly higher for BMP50 than for BG (**Figure 7D**). Furthermore, the amount of bone grown outside of the implant did not depend on the BMP-2 dose (**Figure SI 8**). Finally, the homogeneity score (HS) of bone inside the scaffold was similar for the low and high BMP-2 doses **Figure 7E and Figure SI9**). We also compared the quantitative results of the µCT analysis to those of the CT analysis and the CT-scan score (**Figure SI 10**). The quantification of the new bone volume using μCT acquisitions and the CT-scan score correlated linearly with a regression cofficient R^2^=0.82.

**FIGURE 6.**
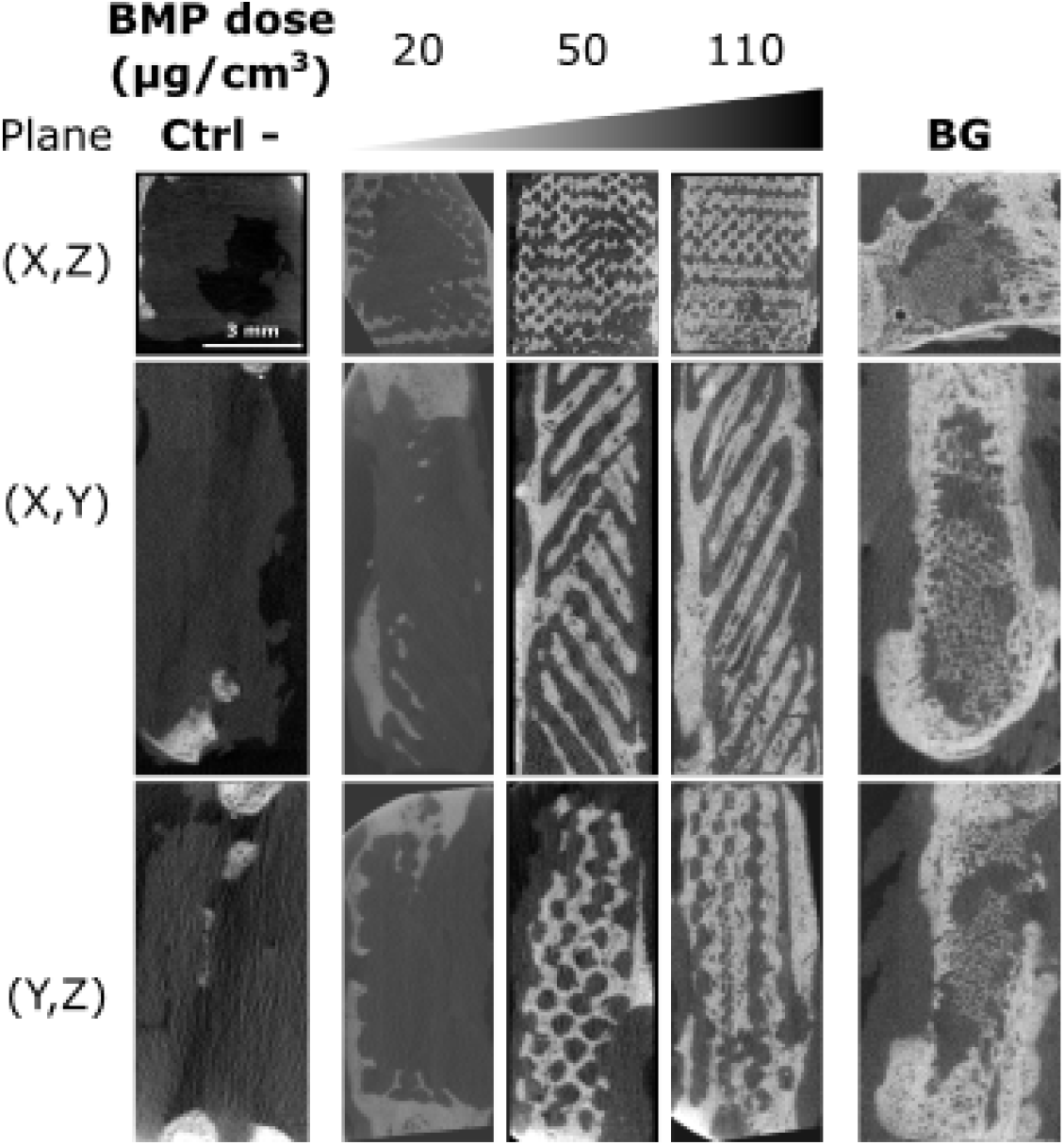
Representative μCT images in the three sectional planes (see Figure 1) for the film-coated scaffolds containing increasing doses of BMP-2 from 20 to 110 μg/cm^3^). The negative control (Ctrl-, film-coated scaffold) is shown on the left, while the bone autograft BG is shown on the right-hand side.

**FIGURE 7.**
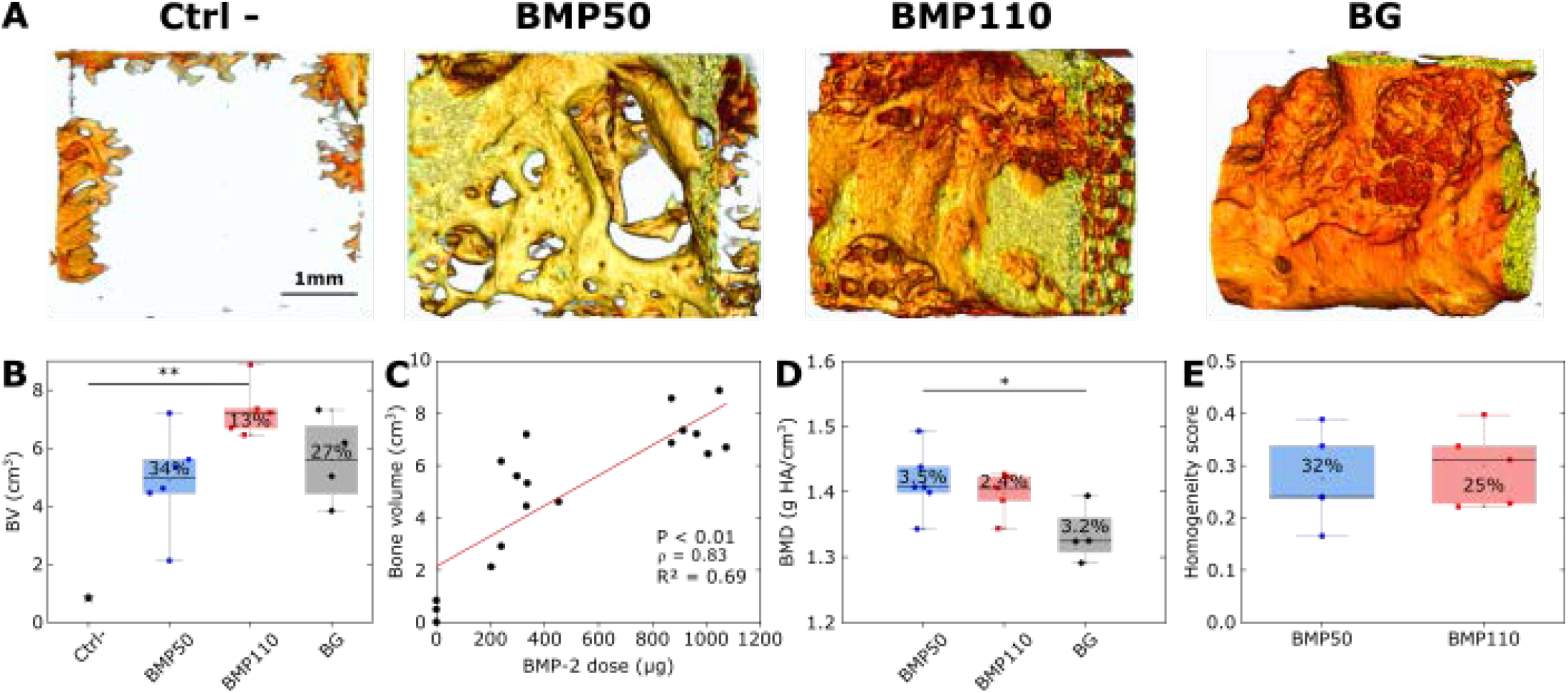
Quantitative μCT analysis of bone formation at 3 months (D91), after explantation. (A) representative images obtained for: the negative control (EDC30 film-coated scaffold without BMP-2), film-coated scaffolds at BMP50 and BMP110 BMP-2 doses, and bone autograft. (B) Box plot representation of the total bone volume as a function of the BMP-2 dose. (C) Total bone volume as a function of the total BMP-2 dose per implant. (D) Bone mineralized density (BMD) as a function of the BMP-2 dose. (E) Homogeneity score measured for BMP50 and BMP110. In each box, the coefficient of variation of the data is given in %. * p < 0.05; ** p < 0.01.

A histological examination (**Figure 8**) revealed that when no BMP-2 was present, only mesenchymal tissue (m) was formed (**Figure 8A**). With BMP50, the amount of new bone was low and mesenchymal tissue was visible (**Figure 8B**). The presence of mature bone with a characteristic Haversian structure in BMP110 implants was evidenced (**Figure 8C**). Imaging under polarized light (**Figure 8D**) allowed a better visualization of the Haversian canals (highlighted by asterisks) and the connections between osteocytes. Furthermore, the interface between the host bone and the newly formed bone was visible (dashed lines in **Figure 8E**), since the host bone had a more lamellar structure than the newly-formed bone (**Figure 8F**). Some bridges were visible between the two types of bones (white arrows in **Figure 8F**), which may contribute to increase the mechanical resistance of the newly formed bone. In some cases, especially for BMP110, the difference between native and new bone was not even distinguishable (data not shown). In the case of BG, host bone and grafted bone were in direct contact (**Figure 8G, H**) or separated by mesenchymal tissue and new bone showed a Haversian structure. The bone area over total area (BA/TA in %) was quantified based on these images (**Figure 8I**). In agreement with the CT and μCT quantifications, more bone was formed for BMP110, whose median value was similar with BG. The homogeneity of the newly formed bone inside the 3D architecture implant was quantified (**Figure 8J**). Here again, bone formation was similar for the three sections of the sample, proving the homogeneity of bone formation. Finally, no local inflammation occurred due to the implant and PLA degradation had not occurred yet (**Figure 8**).

**FIGURE 8:**
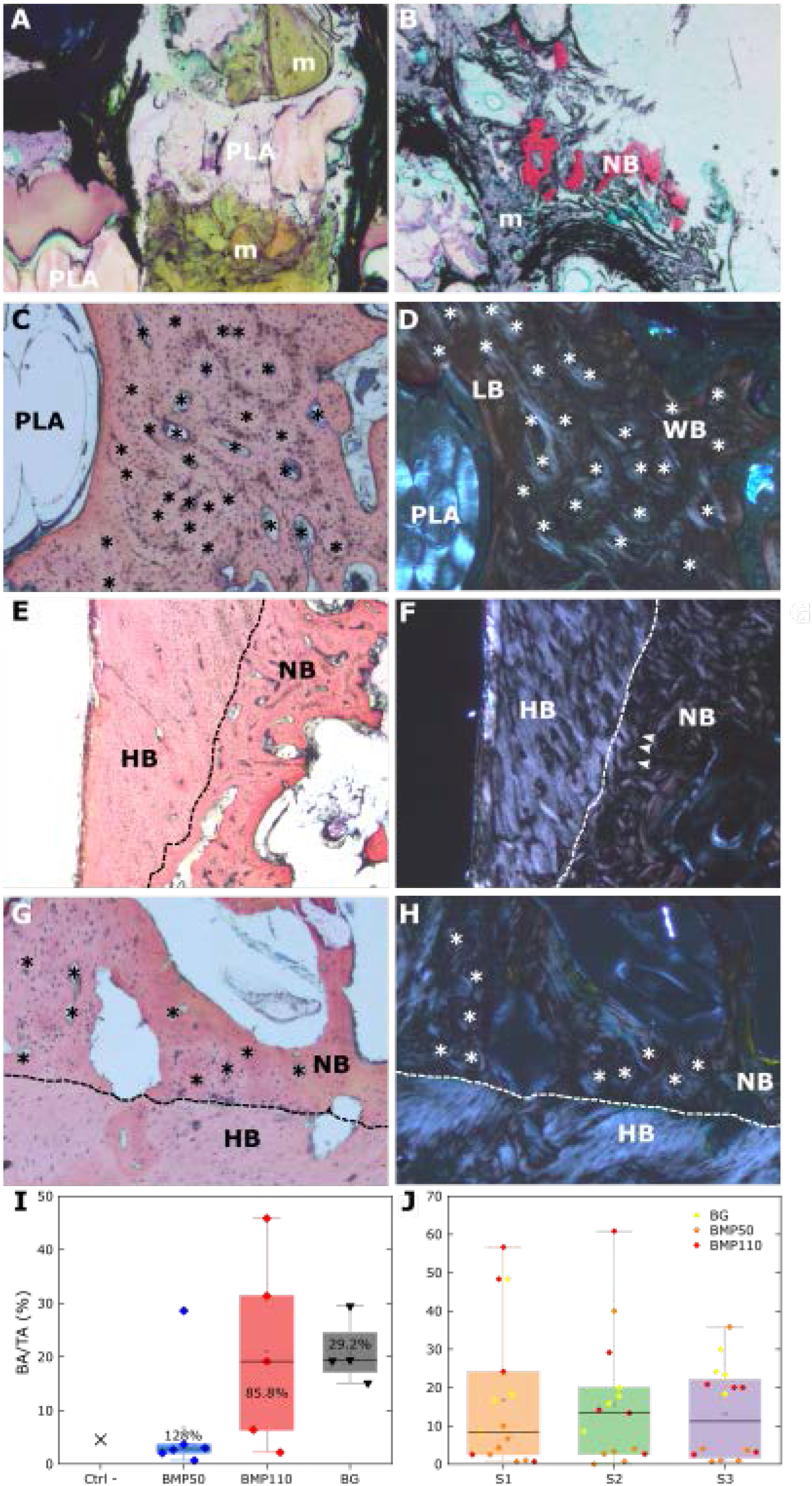
Histological examination and histomorphometry analysis. (A-D) Representative histological sections to show the structure of the newly-formed bone. (A) In the absence of BMP-2 in the film, only mesenchymal tissue (m) is formed (x2 magnification). (B) With low BMP-2 dose (BMP50), new bone is formed along with mesenchymal tissue (x2 magnification). (C, D) The Haversian structure of newly formed bone using BMP-2 at 110 μg/cm^3^ is visible notably in polarized light (x4 magnification). Both lamellar (LB) and woven bones (WB) are visible. Haversian canals are shown with an asterisk (*). (E-H) Representative histological sections to show the contact between host bone and new bone. (E) Contact between host bone and new bone is shown on a transmitted light image of a film-coated implant loaded with high BMP-2 dose (BMP110) highlighted by a dashed line (x2 magnification). (F) The HB/NB interface is clearly visible by polarized light for BMP110, where bony bridges can be observed (white arrows) (x2 magnification). (G, H) Transmitted and polarized light images for BG. The contact between HB and NB as well as the Haversian structure are evidenced (x4 magnification). (I) Quantitative analysis of the bone area over total area (BA/TA) (J) % of bone inside each section of the implant (S1, S2, S3).

Altogether, these data show that bone formation inside the 3D architectured scaffold is spatially homogeneous, and that there is a significant BMP-2 dose-dependent bone formation. BMP-2 mostly influences the formation of mineralized bone and does not induce the formation of ectopic bone.

## Discussion

Before applying new osteogenic technologies in clinical practice, it is necessary to confirm their effectiveness in preclinical studies using critical-size bone defects in relevant animal models. A critical-size bone defect is a defect that does not spontaneously heal for the duration of the study. Our aim was to create a critical-size bone defect close to clinical situations, particularly in maxillofacial surgery where it is frequently needed to fill defects over 10 cm^3^. Among the different animal models published, the pig is a very good candidate since the pig mandible mimics the human mandible in size, anatomy, form, and blood supply [29]. The bone physiology of the pig is very close to human bone physiology, with lamellar and Haversian structure. Several studies have reported on a mandibular defect in adult pigs. There is no real consensus on the critical size, it varied from 2 to 10 cm^3^ depending on the location (tooth bearing area or not, full thickness or not, anterior or posterior area, periosteum preservation or not…) [31]. Otto and coworkers recently proposed a 6 cm^3^ mandibular defect [32]. We carried out a full thickness defect on the basilar border of the mandibular angle. The defect measured 3 x 4 cm, in a portion of mandible that was 1 cm thick, for a total defect volume of 12 cm^3^. To our knowledge, to date, there is no publication dealing with such a volume for a critical-sized bone defect in an animal model [3]. The buccal and vestibular periosteum was removed around the defect to avoid spontaneous ossification, and the age of the animals (24 months) precluded any growth potential as bone maturation had already occurred. The preservation of the alveolar ridge is another advantage of our model. The defect was created far from the teeth, and there was no opening into the oral cavity. This limits the risk for oral fistula and infection. The location of the defect at the mandibular angle permitted a very stable fixation with two reconstructive plates fixed with bicortical locked screws inserted into thick bone. We did not have mechanical complications (material loosening, plate or screw rupture) during the study. The negative control in our study (**Figure 3, 4, and 6**) shows the validity of this critical-size bone defect model.

Long-term biodegradable polymers are interesting for bone tissue engineering applications and 3D printing to create porous scaffolds, since they can provide a temporary mechanical support while leaving sufficient space and time for the newly formed bone to grow. 3D printed scaffolds are particularly suited for the repair of large bone defects. The family of polylactides (PLA, PGA) and polycaprolactone (PCL) are the most widely used for bone tissue engineering [33]. 3D printing of PCL/HAP scaffolds using FDM has already been established [10, 34] [12]. It was shown that the porous scaffolds act as a guiding substrate to enable the formation of a structured fibrous network as a prerequisite for later bone formation [34], the HAP exhibiting osteoconductive properties. Later studies showed that this scaffold can be combined with a rhBMP-7 paste, and that BMP-7 accelerates the bone healing kinetics without changing the bone microstructure [35]. The same team used a critical-size sheep tibial model (30 mm long, 20 mm in diameter, i.e. a total volume of 6 cm^3^ with a central empty space of 2.4 cm^3^) to investigate whether a BMP-7 paste added inside the scaffold at two doses (1.75 and 3.5 mg) could influence bone growth. They did not see any BMP-7 dose-dependence [12]. As an alternative we used PLA that is also widely employed in orthopedic and maxillofacial surgery [7] due to its long-term biodegradability and modularity [36]. It also has interesting mechanical properties compared to PCL [7]. For the first time to our knowledge, we developed a 3D printed PLA implant of clinical grade using FDM. This fabrication process allowed for a complete control over the scaffold architecture. Indeed, compared to other widely-used manufacturing methods such as freeze-drying for example, the porosity can be controlled as well as the external shape of the scaffold [37]. Our previous studies had shown that the polyelectrolyte film can be deposited on PLGA cylinders [21] and PLA microcarriers [38], which was confirmed here on 3D printed scaffolds. The PLA being inert, it does not provide osteoinduction and the osteoinductive properties are exclusively provided via the BMP-2-containing film coating. Thanks to the homogeneous surface coating of the 3D printed scaffold by the film, bone formation was highly homogeneous inside all the pores of the scaffold (**Figure 6-8**), and there was no general or local inflammatory reaction (**Figure 8 and Figure SI 1**). In additon, the PLA degrades over long term, typically from 6 months to 2 years [39] and no sign of degradation was found in the histological images (**Figure 8**).

BMP-2 dose is a critical issue for clinical applications. It has been demonstrated that the osteogenic response is correlated to the dose of BMP-2 [40]. However, several studies showed that high doses are correlated with adverse effects [16, 21], but it is difficult to find a consensus in the literature regarding the effective dose of BMP-2 to achieve the expected clinical results. Boden et al. [41] mentioned in their publication following a randomized clinical pilot trial for posterolateral lumbar fusion that the required dose (i.e. the concentration of growth factor expressed per volume of final collagen carrier matrix) to induce 100% consistent bone formation was substantially higher in non-human primates (1.5 - 2.0 mg/mL) than in rodents (0.2 - 0.4 mg/mL). James et al [17] reviewed the side effects of BMP-2 and noted that excessive concentrations of BMP-2 may be the most important factor contributing to the majority of adverse events: they noticed that increasing doses of BMP-2 do not necessarily result in higher fusion rates in spine procedures and long bone non-unions. These complications, described in clinical trials or in experimental studies, included postoperative inflammation (from benign seroma to life-threatening effects), ectopic bone formation, bone resorption and bone cyst formation, urogenital events (retrograde ejaculation, bladder retention) and wound complications [17, 42, 43]. The implication of BMP-2 in carcinogenesis is controversial. For all these reasons, it is important to reduce the dose of BMP-2 and to deliver it locally and progressively. Here, we considerably reduced the amount of BMP-2 with a dose of 240 to 1000 μg per 12 cm^3^ defect, which corresponded to 0.02 to 0.08 mg/cm^3^ (1 mg/cm^3^ is equivalent to mg/mL). This corresponds to a 20-fold to 75-fold reduction compared to the commercially available collagen sponges, which are approved for a dose of 1.5 mg/mL. We chose these BMP-2 doses following a literature analysis, indicating that the studies were previously conducted in the range from 0.03 to 3 mg/cm^3^ [16, 44-46] [47] and based on our previous studies in rat and rabbit [21, 48]. Our aim was to reduce the BMP-2 dose compared to what was usually done.

Regarding the difference between EDC30 and EDC70 films in the BMP-2 dose-dependent bone regeneration (**Fig. 3**), we may hypothesize that it may be due to several reasons: i) the different stiffness of the EDC30 and EDC70 films, ii) the different loading of BMP-2 in the EDC30 and EDC70 films (Fig 2D), the amount of BMP-2 loaded being higher for the EDC30 films at high BMP-2 concentrations, iii) the higher release for the EDC30 films as shown in Figure 2E. iv) the fact that BMP-2 internalization by cells is higher for EDC30 than EDC70 [49].

We did not notice adverse effects such as local inflammation, swelling, bone cysts or bone resorption. In three cases, we noticed during the explantation a small seroma of less than 1 mL: two cases for the bone autograft and one case for high dose BMP-2. The amount of ossification outside the scaffold was small and independent of the BMP-2 dose (**Figure 6**).

The visual scoring used by clinicians for the analysis of CT-scans is a fine and simple way to evaluate bone regeneration. While it can be considered as subjective, the number of independent operators (four in our case) and the fact that this is done in a blind manner prevent the risk of a biased view. Furthermore, the results correlated well to to the quantitative analysis (**Figure SI 10**). Another reason for using this score was to be close to the kind of evaluations that are usually conducted in the clinical routine.

Similarly, the histomorphometry was conducted in a blind manner by three independent operators (**Figure 8I and 8J**). Though this may result in over-estimation of the amount of new bone formation, it remains a simple and effective way to compare different conditions. Here, a systematic and purely objective evaluation was not possible. Again, the results were in accordance with the previous quantitative analyses (**Figure 5 and 6**).

Our study opens perspectives for the clinical translation of these osteoinductive 3D printed PLA scaffolds. First, the PLA used was clinical-grade and the film components are already approved by the US Food and Drug Administration (FDA) and the European Medicine Agencies (EMA) [21]. 3D printing is a cost-effective solution [10]. It has the potential to produce new medical products with unprecedented structural and functional designs, and its regulatory landscape is rapidly evolving [9]. The fact that the new synthetic graft proposed here is made of a 3D printed scaffold and a 2D osteoinductive film coating containing BMP-2 offers several modularity possibilities in terms of scaffold design (dimensions, shape, architecture, porosity) and a precise control of BMP-2 dose delivered via the film coating. Regulations regarding the osteoinductive biomolecule, the BMP-2 growth factor, is also less difficult than that of stem cells [9], and is already clinically approved for several indications.

## Conclusions

In summary, we engineered a 3D printed scaffold made by FDM and coated with a biomimetic polyelectrolyte film loaded with BMP-2 to repair a new model of critical-size bone defect in mini-pig mandible. This bone defect model and volume > 10 cm^3^ is equivalent to various clinical applications in humans, showing the translation potential of our technology. The 3D architecture of the scaffold provided a guide for cells to grow inside the volumetric defect while the 2D osteoinductive coating allowed BMP-2 to trigger cell differentiation and bone regeneration. We showed that the BMP-2 dose delivered from the polyelectrolyte film significantly influenced the amount and maturity of regenerated bone with a clear BMP-2 dose-dependence for EDC30. The repair kinetics was also dose-dependent, with a slower kinetics for the high BMP-2 dose. CT scans, μCT acquisitions and histological examinations proved the formation of mineralized and well-vascularized bone with high BMP-2 doses while lower BMP-2 doses lead to less mineralized bone. In addition, the new bone formed homogeneously inside the scaffold, whatever the BMP-2 dose, and with little ectopic bone formation. This combination of a 3D printed scaffold with a 2D osteoinductive coating opens perspectives in personalized medicine since 3D printing allows the customization of shape of implants and the biomimetic coating allows the controlled delivery of BMP-2 in space and time.

## Supplementary Materials

Figure SI 1. Analysis of blood markers.

Figure SI 2. CT scans of the samples of the preliminary experiment (EDC30 & EDC70, two BMP-2 doses) and negative control.

Figure SI 3. Quantitative analysis of poorly and highly mineralized bone volume over time using CT images. Figure

SI 4. Micro-computed tomography images of the preliminary experiment: bone volume and bone mineral density.

Figure SI 5. Micro-computed tomography images in the three planes of BMP-2 loaded films (EDC70). Figure SI 6. Histological analysis of the preliminary experiment. Quantification of bone area over total area (BA/TA).

Figure SI 7. 3D reconstructed images of scaffolds as a function of the BMP-2 dose loaded in the scaffold. Figure

SI 8. Bone volume growing outside of the implant for BMP50 and BMP110.

Figure SI 9. Analysis of bone homogeneity within the scaffold (homogeneity score).

Figure SI 10. Correlation between the CT-scan score and the quantitative analyses of new bone volume formed.

## Acknowledgments

We thank the following for their contribution: Isabelle Paintrand (CNRS), Jie Liu (Grenoble Institute of Technology) for help in the preparation of the samples; Jean-Luc Coll (Institute of Advanced Biosciences) and Remy Gerez (Institut Claude Bourgelat) for fruitful discussions, Heiko Richter and Birgitta Stolze (LLS Rowiak) for technical discussions, R Lartizien (Annecy Genenevois hospital) for help in the clinical score, Sebastien Schoumacker for help in 3D printing, Dorothée Palluy (Stryker) for providing the reconstruction plates and screws, Sylvie Berthier (CHU-Grenoble Alpes) for her help in the quantitative histological analysis.

## CRediT author contribution statement: Michaël Bouyer

Conceptualization, Formal analysis, Investigation, Methodology, Visualization. **Charlotte Garot:** Formal analysis, Writing – original draft, Writing – review and editing, Visualization. **Paul Machillot:** Investigation. **Julien Vollaire:** Formal analysis, Investigation, Writing – original draft. **Vincent Fitzpatrick:** Investigation. **Sanela Morand:** Formal analysis. **Jean Boutonnat:** Formal analysis. **Véronique Josserand:** Formal analysis. **Georges Bettega:** Conceptualization, Investigation, Methodology, Writing – original draft, Writing – review and editing, Supervision. **Catherine Picart:** Conceptualization, Methodology, Writing – original draft, Writing – review and editing, Supervision, Funding acquisition.

## Data availability

The raw/processed data required to reproduce these findings cannot be shared at this time due to legal reasons. They are currently available upon request and will be deposited in a data repository.

## Bibliographic references

[1] C. Laurencin, Y. Khan, S.F. El-Amin, Bone graft substitutes, Expert review of medical devices 3(1) (2006) 49–57.

[2] M. Woodruff, C. Lange, J. Reichert, A. Berner, F. Chen, P. Fratzl, J.J.-T. Schantz, D. Hutmacher, Bone tissue engineering: from bench to bedside, Materials Today 15(10) (2012) 430–432.

[3] S. Zeiter, K. Koschitzki, M. Alini, F. Jakob, M. Rudert, M. Herrmann, Evaluation of preclinical models for the testing of bone tissue-engineered constructs, Tissue Eng Part C Methods (2020).

[4] A. Oryan, S. Alidadi, A. Moshiri, N. Maffulli, Bone regenerative medicine: classic options, novel strategies, and future directions, Journal of orthopaedic surgery and research 9(1) (2014) 18.

[5] J.C. Reichert, A. Cipitria, D.R. Epari, S. Saifzadeh, P. Krishnakanth, A. Berner, M.A. Woodruff, H. Schell, M. Mehta, M.A. Schuetz, G.N. Duda, D.W. Hutmacher, A tissue engineering solution for segmental defect regeneration in load-bearing long bones, Sci Transl Med 4(141) (2012) 141ra93.

[6] X.P. Tan, Y.J. Tan, C.S.L. Chow, S.B. Tor, W.Y. Yeong, Metallic powder-bed based 3D printing of cellular scaffolds for orthopaedic implants: A state-of-the-art review on manufacturing, topological design, mechanical properties and biocompatibility, Mater Sci Eng C Mater Biol Appl 76 (2017) 1328–1343.

[7] Z. Sheikh, S. Najeeb, Z. Khurshid, V. Verma, H. Rashid, M. Glogauer, Biodegradable materials for bone repair and tissue engineering applications, Materials 8(9) (2015) 5744–5794.

[8] A. Cheng, Z. Schwartz, A. Kahn, X. Li, Z. Shao, M. Sun, Y. Ao, B.D. Boyan, H. Chen, Advances in porous scaffold design for bone and cartilage tissue engineering and regeneration, Tissue Eng Part B Rev 25(1) (2019) 14–29.

[9] L.M. Ricles, J.C. Coburn, M. Di Prima, S.S. Oh, Regulating 3D-printed medical products, Sci Transl Med 10(461) (2018).

[10] A. Youssef, S.J. Hollister, P.D. Dalton, Additive manufacturing of polymer melts for implantable medical devices and scaffolds, Biofabrication 9(1) (2017) 012002.

[11] X. Liang, J. Gao, W. Xu, X. Wang, Y. Shen, J. Tang, S. Cui, X. Yang, Q. Liu, L. Yu, J. Ding,Structural mechanics of 3D-printed poly(lactic acid) scaffolds with tetragonal, hexagonal and wheel-like designs, Biofabrication 11(3) (2019) 035009.

[12] A. Cipitria, J.C. Reichert, D.R. Epari, S. Saifzadeh, A. Berner, H. Schell, M. Mehta, M.A. Schuetz, G.N. Duda, D.W. Hutmacher, Polycaprolactone scaffold and reduced rhBMP-7 dose for the regeneration of critical-sized defects in sheep tibiae, Biomaterials 34(38) (2013) 9960–8.

[13] M.A. Brennan, A. Renaud, J. Amiaud, M.T. Rojewski, H. Schrezenmeier, D. Heymann, V. Trichet, P. Layrolle, Pre-clinical studies of bone regeneration with human bone marrow stromal cells and biphasic calcium phosphate, Stem Cell Res Ther 5(5) (2014) 114.

[14] M.R. Urist, Bone -formation by autoinduction, Science 150(3698) (1965) 893-&.

[15] V.E. Santo, M.E. Gomes, J.F. Mano, R.L. Reis, Controlled release strategies for bone, cartilage, and osteochondral engineering--Part II: challenges on the evolution from single to multiple bioactive factor delivery, Tissue Eng Part B Rev 19(4) (2013) 327–52.

[16] J.N. Zara, R.K. Siu, X. Zhang, J. Shen, R. Ngo, M. Lee, W. Li, M. Chiang, J. Chung, J. Kwak, B.M. Wu, K. Ting, C. Soo, High doses of bone morphogenetic protein 2 induce structurally abnormal bone and inflammation in vivo, Tissue Eng Part A 17(9-10) (2011) 1389–99.

[17] A.W. James, G. LaChaud, J. Shen, G. Asatrian, V. Nguyen, X. Zhang, K. Ting, C. Soo, A review of the clinical side effects of bone morphogenetic protein-2, Tissue Eng Part B Rev 22(4) (2016) 284–97.

[18] H.J. Seeherman, S.P. Berasi, C.T. Brown, R.X. Martinez, Z.S. Juo, S. Jelinsky, M.J. Cain, J. Grode, K.E. Tumelty, M. Bohner, O. Grinberg, N. Orr, O. Shoseyov, J. Eyckmans, C. Chen, P.R. Morales, C.G. Wilson, E.J. Vanderploeg, J.M. Wozney, A BMP/activin A chimera is superior to native BMPs and induces bone repair in nonhuman primates when delivered in a composite matrix, Sci Transl Med 11(489) (2019).

[19] W.J. King, P.H. Krebsbach, Growth factor delivery: how surface interactions modulate release in vitro and in vivo, Adv. Drug. Deliv. Rev. 64(12) (2012) 1239–56.

[20] E. Migliorini, A. Valat, C. Picart, E.A. Cavalcanti-Adam, Tuning cellular responses to BMP-2 with material surfaces, Cytokine Growth Factor Rev 27 (2016) 43–54.

[21] M. Bouyer, R. Guillot, J. Lavaud, C. Plettinx, C. Olivier, V. Curry, J. Boutonnat, J.L. Coll, F. Peyrin, V. Josserand, G. Bettega, C. Picart, Surface delivery of tunable doses of BMP-2 from an adaptable polymeric scaffold induces volumetric bone regeneration, Biomaterials 104 (2016) 168–81.

[22] J.A. McGovern, M. Griffin, D.W. Hutmacher, Animal models for bone tissue engineering and modelling disease, Disease models & mechanisms 11(4) (2018).

[23] T. Crouzier, L. Fourel, T. Boudou, C. Albiges-Rizo, C. Picart, Presentation of BMP-2 from a soft biopolymeric film unveils its activity on cell adhesion and migration, Adv. Mater. 23(12) (2011) H111–8.

[24] T. Crouzier, K. Ren, C. Nicolas, C. Roy, C. Picart, Layer-by-Layer films as a biomimetic reservoir for rhBMP-2 delivery: controlled differentiation of myoblasts to osteoblasts, Small 5(5) (2009) 598–608.

[25] R. Guillot, F. Gilde, P. Becquart, F. Sailhan, A. Lapeyrere, D. Logeart-Avramoglou, C. Picart, The stability of BMP loaded polyelectrolyte multilayer coatings on titanium, Biomaterials 34(23) (2013) 5737–46.

[26] T. Crouzier, F. Sailhan, P. Becquart, R. Guillot, D. Logeart-Avramoglou, C. Picart, The performance of BMP-2 loaded TCP/HAP porous ceramics with a polyelectrolyte multilayer film coating, Biomaterials 32(30) (2011) 7543–54.

[27] C. Kunert-Keil, H. Richter, I. Zeidler-Rentzsch, I. Bleeker, T. Gredes, Histological comparison between laser microtome sections and ground specimens of implant-containing tissues, Annals of Anatomy 222 (2019) 153–157.

[28] J.K. Stembirek, M Putnova, I ; Stehlik, L ; Buchtova, M The pig as an experimental model for clinical craniofacial research, Laboratory Animals 2012; 46: 269–279 46 (2012) 269–279.

[29] B. Saka, A. Wree, L. Anders, K.K. Gundlach, Experimental and comparative study of the blood supply to the mandibular cortex in Gottingen minipigs and in man, J Craniomaxillofac Surg 30(4) (2002) 219–25.

[30] Z. Sun, K.S. Kennedy, B.C. Tee, J.B. Damron, M.J. Allen, Establishing a critical-size mandibular defect model in growing pigs: characterization of spontaneous healing, J Oral Maxillofac Surg 72(9) (2014) 1852–68.

[31] J.L. Ma, J.L. Pan, B.S. Tan, F.Z. Cui, Determination of critical size defect of minipig mandible, J Tissue Eng Regen Med 3(8) (2009) 615–22.

[32] F.A. Probst, R. Fliefel, E. Burian, M. Probst, M. Eddicks, M. Cornelsen, C. Riedl, H. Seitz, A. Aszodi, M. Schieker, S. Otto, Bone regeneration of minipig mandibular defect by adipose derived mesenchymal stem cells seeded tri-calcium phosphate-poly(D,L-lactide-co-glycolide) scaffolds, Sci Rep 10(1) (2020) 2062.

[33] D.W. Hutmacher, Scaffolds in tissue engineering bone and cartilage, Biomaterials 21(24) (2000) 2529–43.

[34] A. Cipitria, C. Lange, H. Schell, W. Wagermaier, J.C. Reichert, D.W. Hutmacher, P. Fratzl, G.N. Duda, Porous scaffold architecture guides tissue formation, J Bone Miner Res 27(6) (2012) 1275–88.

[35] A. Cipitria, W. Wagermaier, P. Zaslansky, H. Schell, J.C. Reichert, P. Fratzl, D.W. Hutmacher, G.N. Duda, BMP delivery complements the guiding effect of scaffold architecture without altering bone microstructure in critical-sized long bone defects: A multiscale analysis, Acta Biomater. 23 (2015) 282–294.

[36] J.E. Bergsma, W.C. de Bruijn, F.R. Rozema, R.R. Bos, G. Boering, Late degradation tissue response to poly(L-lactide) bone plates and screws, Biomaterials 16(1) (1995) 25–31.

[37] C. Garot, G. Bettega, C. Picart, Additive manufacturing of material Scaffolds for bone regeneration: toward application in the clinics, Adv. Funct. Mat. n/a(n/a) (2020) 2006967.

[38] M. Kuterbekov, P. Machillot, P. Lhuissier, C. Picart, A.M. Jonas, K. Glinel, Solvent-free preparation of porous poly(l-lactide) microcarriers for cell culture, Acta Biomater. 75 (2018) 300–311.

[39] S. Farah, D.G. Anderson, R. Langer, Physical and mechanical properties of PLA, and their functions in widespread applications - A comprehensive review, Adv. Drug. Deliv. Rev. 107 (2016) 367–392.

[40] J.D. Boerckel, Y.M. Kolambkar, K.M. Dupont, B.A. Uhrig, E.A. Phelps, H.Y. Stevens, A.J. Garcia, R.E. Guldberg, Effects of protein dose and delivery system on BMP-mediated bone regeneration, Biomaterials 32(22) (2011) 5241–51.

[41] S.D. Boden, J. Kang, H. Sandhu, J.G. Heller, Use of recombinant human bone morphogenetic protein-2 to achieve posterolateral lumbar spine fusion in humans: a prospective, randomized clinical pilot trial: 2002 Volvo Award in clinical studies, Spine (Phila Pa 1976) 27(23) (2002) 2662–73.

[42] A.F. Kamal, O.S.H. Siahaan, J. Fiolin, Various dosages of BMP-2 for management of massive bone defect in Sprague Dawley rat, Arch Bone Jt Surg 7(6) (2019) 498–505.

[43] E.L. Durham, R.N. Howie, S. Hall, N. Larson, B. Oakes, R. Houck, Z. Grey, M. Steed, A.C. LaRue, R. Muise-Helmericks, J. Cray, Optimizing bone wound healing using BMP2 with absorbable collagen sponge and Talymed nanofiber scaffold, J Transl Med 16(1) (2018) 321.

[44] P. Sun, J. Wang, Y. Zheng, Y. Fan, Z. Gu, BMP2/7 heterodimer is a stronger inducer of bone regeneration in peri-implant bone defects model than BMP2 or BMP7 homodimer, Dent Mater J 31(2) (2012) 239–48.

[45] J. Wang, Y. Zheng, J. Zhao, T. Liu, L. Gao, Z. Gu, G. Wu, Low-dose rhBMP2/7 heterodimer to reconstruct peri-implant bone defects: a micro-CT evaluation, J Clin Periodontol 39(1) (2012) 98–105.

[46] M.H. Pelletier, R.A. Oliver, C. Christou, Y. Yu, N. Bertollo, H. Irie, W.R. Walsh, Lumbar spinal fusion with beta-TCP granules and variable Escherichia coli-derived rhBMP-2 dose, Spine J 14(8) (2014) 1758–68.

[47] M. Hoshino, T. Egi, H. Terai, T. Namikawa, K. Takaoka, Repair of long intercalated rib defects using porous beta-tricalcium phosphate cylinders containing recombinant human bone morphogenetic protein-2 in dogs, Biomaterials 27(28) (2006) 4934–40.

[48] R. Guillot, I. Pignot-Paintrand, J. Lavaud, A. Decambron, E. Bourgeois, V. Josserand, D. Logeart-Avramoglou, E. Viguier, C. Picart, Assessment of a polyelectrolyte multilayer film coating loaded with BMP-2 on titanium and PEEK implants in the rabbit femoral condyle, Acta Biomater. (2016).

[49] F. Gilde, L. Fourel, R. Guillot, I. Pignot-Paintrand, T. Okada, V. Fitzpatrick, T. Boudou, C. Albiges-Rizo, C. Picart, Stiffness-dependent cellular internalization of matrix-bound BMP-2 and its relation to Smad and non-Smad signaling, Acta Biomater. 46 (2016) 55–67.

