## Supplemental File for "3D printed scaffold combined to 2D osteoinductive coatings to repair a critical-size mandibular bone defect"

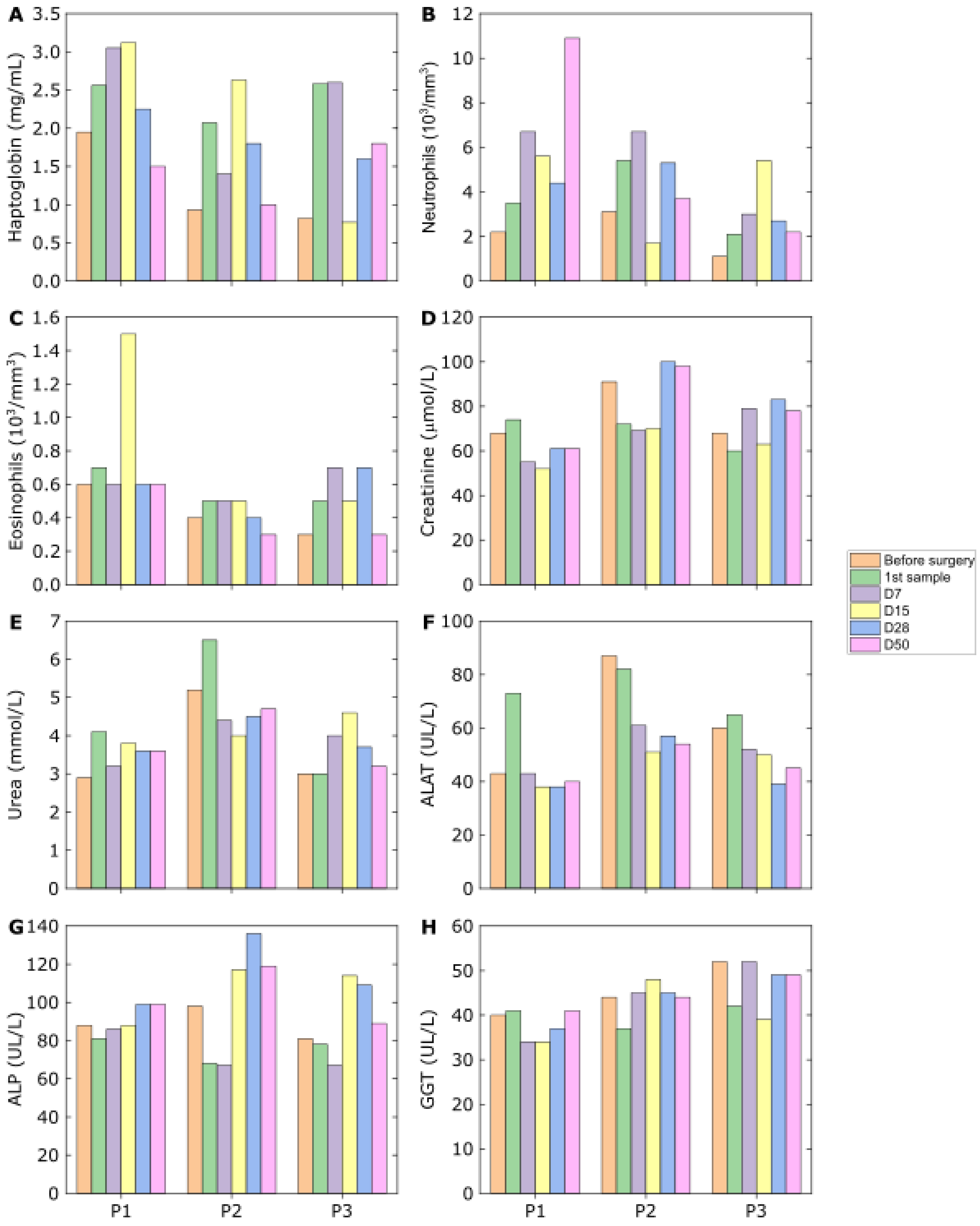


**Figure SI 1**. Blood sample analysis for three different pigs (P1, P2, P3) from the preliminary experiment. P1 was implanted with BMP20 scaffolds (one EDC30 and one EDC70), P2 with BMP110 scaffolds (one EDC30 and one EDC70), and P3 with negative controls (one film-coated scaffold at EDC70 and one empty defect). Several markers were studied at different time points until day 50: (A) haptoglobin; (B) neutrophils; (C) eosinophils (D) creatinine; (E) urea; (F) alanine aminotransferases (ALAT); (G) alkaline phosphatase (ALP); (H) gamma-glutamine transferase (GGT).

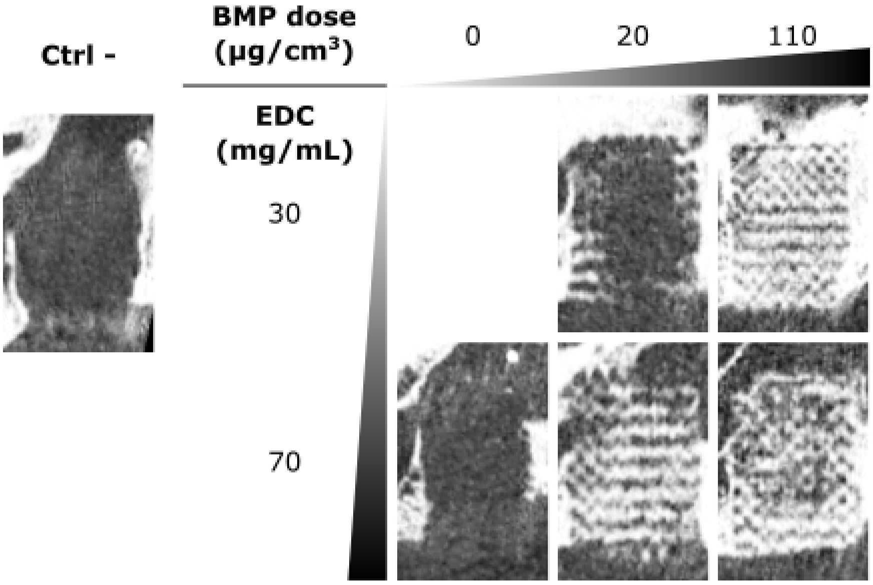


**Figure SI 2.** Computed tomography (CT) imaging at day 91 for the mini-pigs of the preliminary experiment with two film crosslinking levels (EDC30 and EDC70) and two BMP-2 doses (20 and 110 μg/cm$^3$). Two negative controls are presented: one empty defect on the left and one film-coated implant without BMP-2. No BMP-2 dose-dependence was observed for the BMP-2 loaded EDC70 films while a clear dose effect was visible for the EDC30 films.

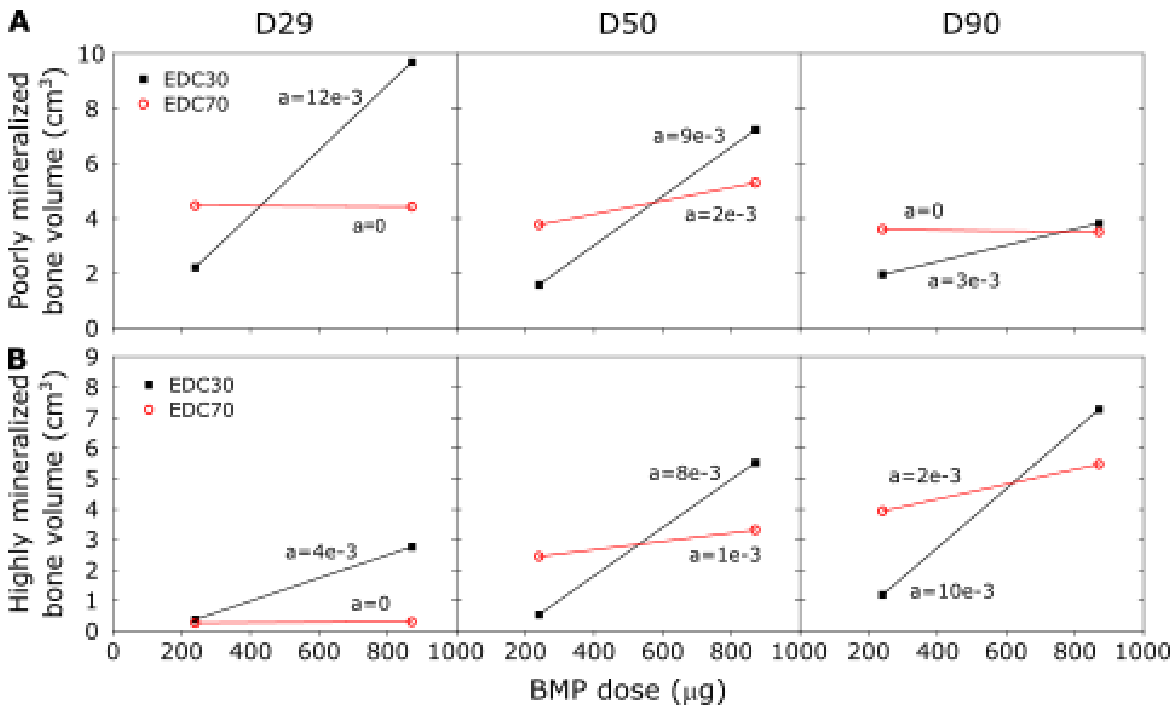


**Figure SI 3**. Quantification of bone volume from the CT images after defining a global threshold: (A) 230 to 629 HU for poorly mineralized bone and (B) > 630 HU for highly mineralized bone. Bone volume is given in $cm^3$. The data corresponding to each film are given in color: black for EDC30 films and red for EDC70 films. The slope is also given. Here again, a dose-dependence is visible for the EDC30 films while there is no systematic change for the EDC70 films.

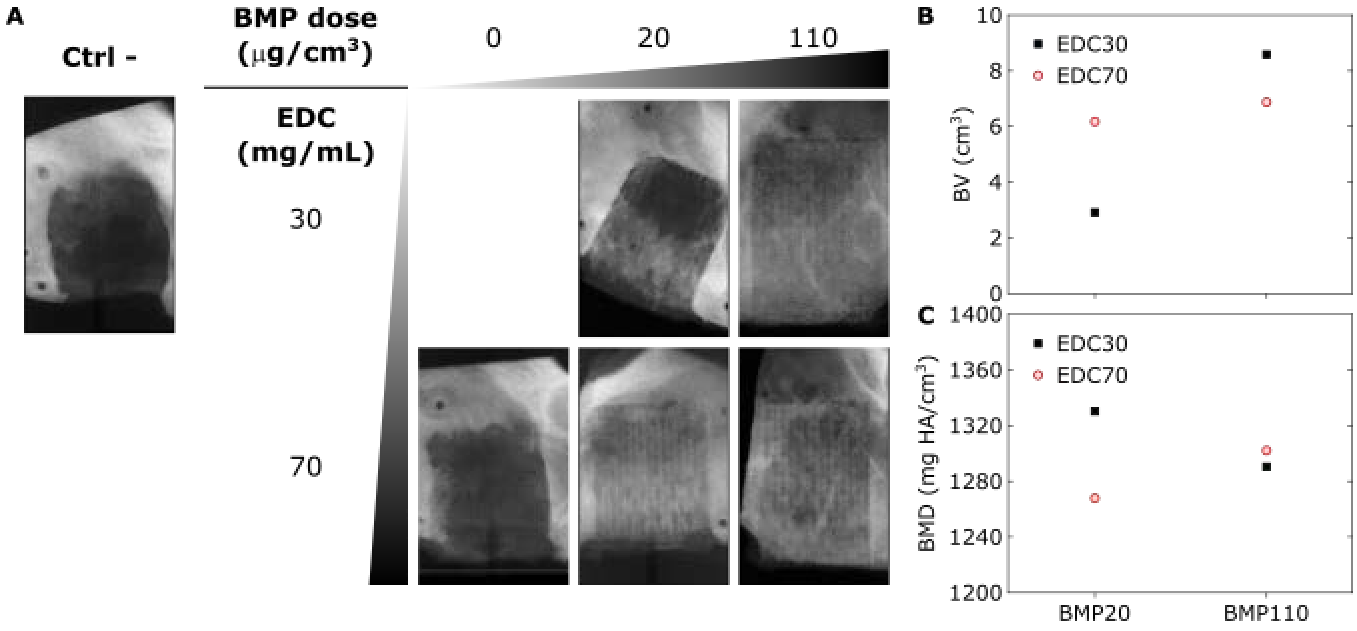


**Figure SI 4**. μCT data acquired after explantation of the scaffolds, for the preliminary experiment. (A) μCT scans with two film crosslinking levels (EDC30 and EDC70) and two BMP-2 doses (20 and 110 μg/cm$^3$). Two negative controls are presented: one empty defect on the left and one film-coated implant without BMP-2. (B) Total bone volume (BV) as a function of the BMP-2 dose and film crosslinking level. (C) Bone mineral density (BMD) as a function of the BMP-2 dose and film crosslinking level. Once again, the BMP-2 dose-dependence is visible with EDC30 but absent with EDC70.

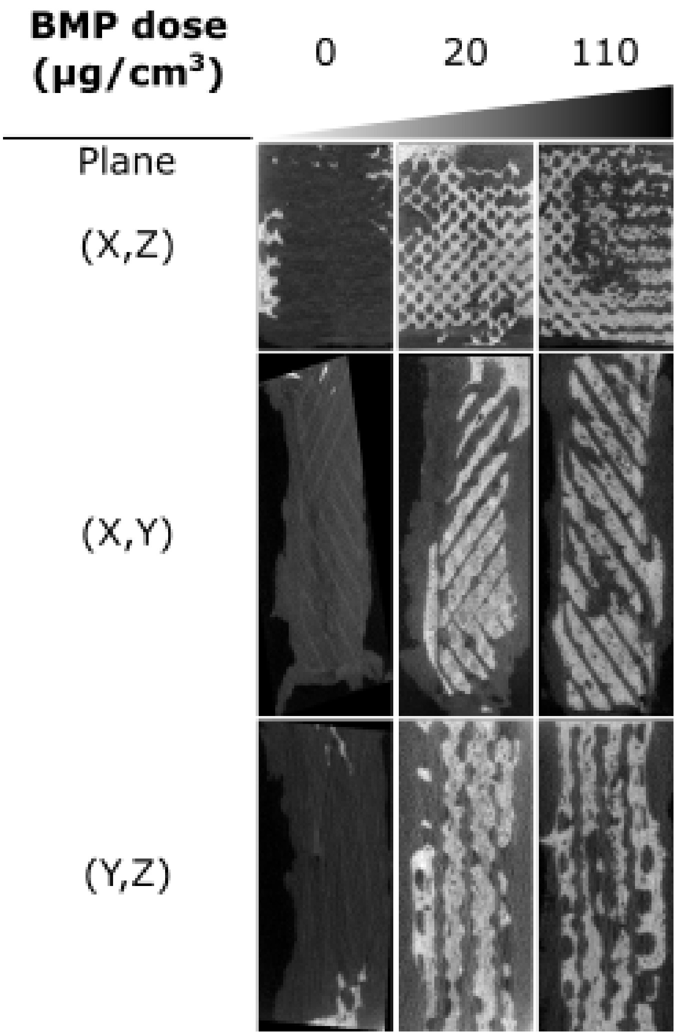


**Figure SI 5.** µCT imaging of the EDC70 film-coated implants given in the three different planes (X,Z), (X,Y), (Y, Z) for the two BMP-2 doses and negative control from the preliminary experiment. Here again, the images were similar, highlighting the lack of BMP-2 dose-dependence for the EDC70 films.

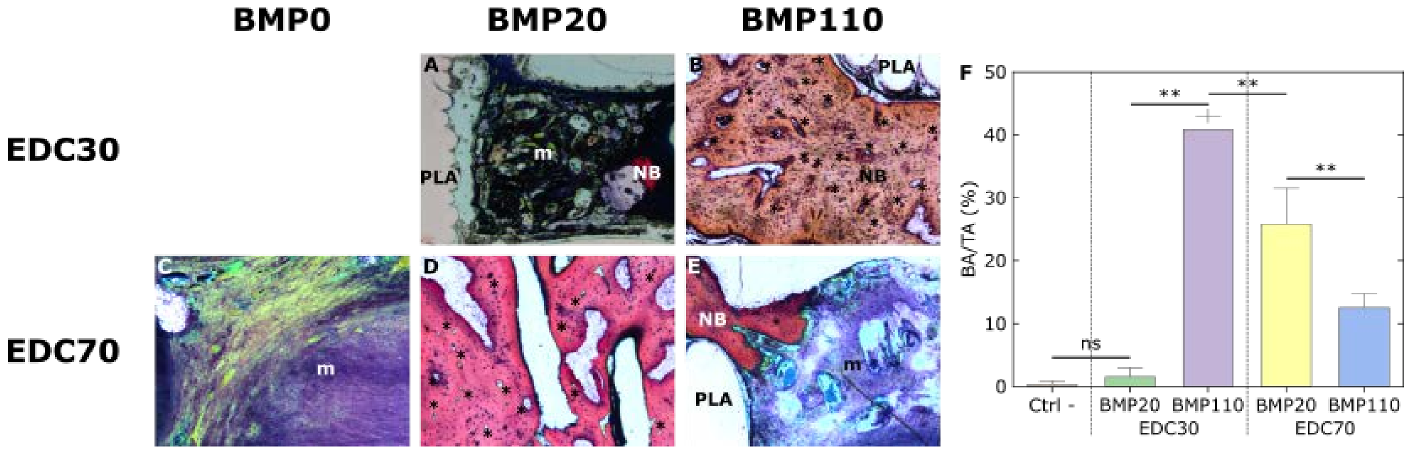


**Figure SI 6.** Histological examination for the samples of the preliminary experiment (x2 magnification for all the images). (A) Representative section of the EDC30 film-coated implant loaded with a low BMP-2 dose (20 µg/cm$^3$). Very few new bone (NB) was formed and mesenchymal tissue (m) was largely observed. (B) EDC30 film-coated implant loaded with a high BMP-2 dose (110 µg/cm$^3$). New bone formation was visible as well as Haversian canals (highlighted by asterisks *). (C) When the scaffold was coated with a film at EDC70 without BMP-2, only mesenchymal tissue (m) was observed. (D) EDC70 film-coated scaffold loaded with BMP20. Haversian canals are visible. (E) EDC70 film-coated scaffold loaded with BMP110. New bone as well as residual mesenchymal tissue can be observed. (F) Quantification of bone area over total area (BA/TA in %) for the different conditions. p** < 0.01. Here again, the BMP-2 dose-dependence of EDC30 films is shown while EDC70 films have a different behavior.

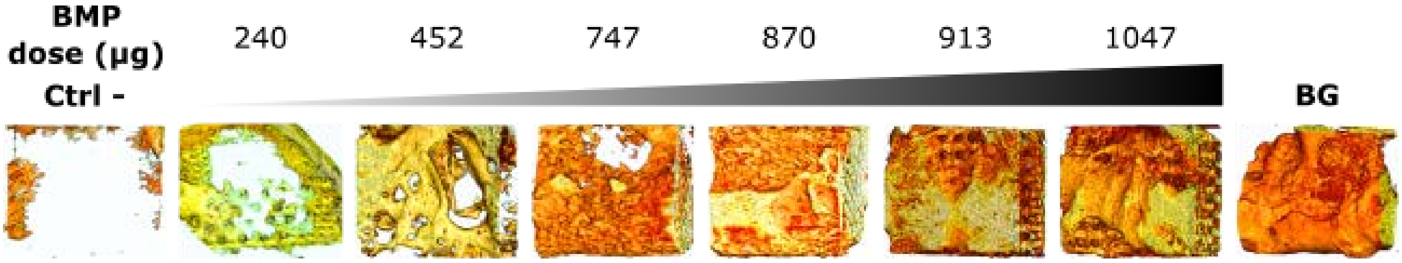


**Figure SI 7**. 3D reconstructed μCT images of individual samples ranged from lowest to highest dose. For each implant, the value of the initial BMP-2 dose loaded is given in μg/implant. The negative control (EDC30 film without BMP-2) and the positive control (bone graft, BG) are also shown.

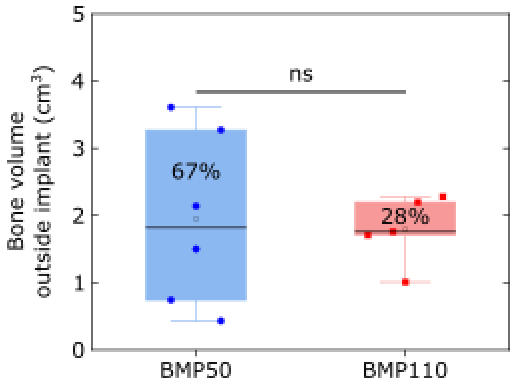


**Figure SI 8**. Bone volume grown outside of the implant, also called “ectopic bone” for the low and high BMP-2 doses (BMP50 and BMP110, respectively).

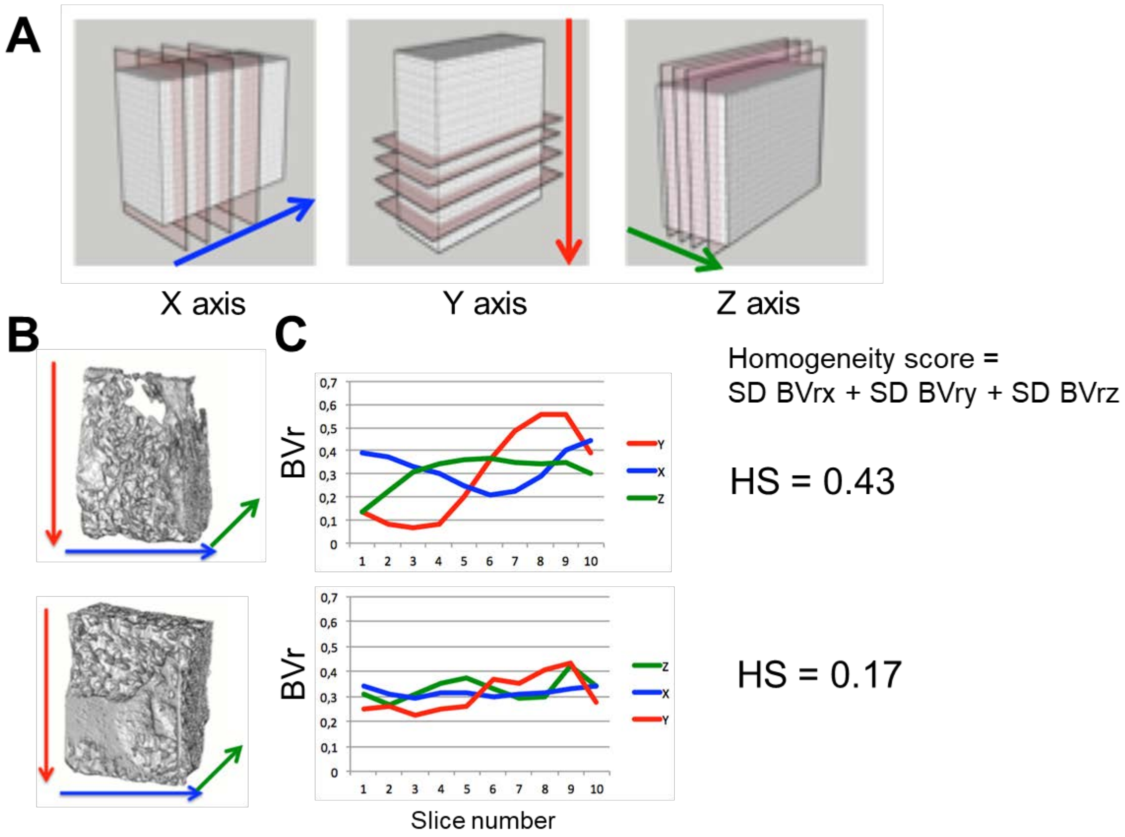


**Figure SI 9**. Calculation of the homogeneity score (HS) for bone grown inside the 3D scaffold. The three axis (X, Y, Z) were considered. The implant was sectionned in 10 slices in each plane along a transverse direction X, Y or Z. The bone volume in each slice was calculated. HS was calculated as the sum of the three standard deviations (SD on BVrx, BVry and BVrz). (B) 3D μCT reconstructed images are shown and the corresponding HS score profiles for two selected bone samples are given here as examples: (B) HS of 0.43 and (C) HS of 0.17.

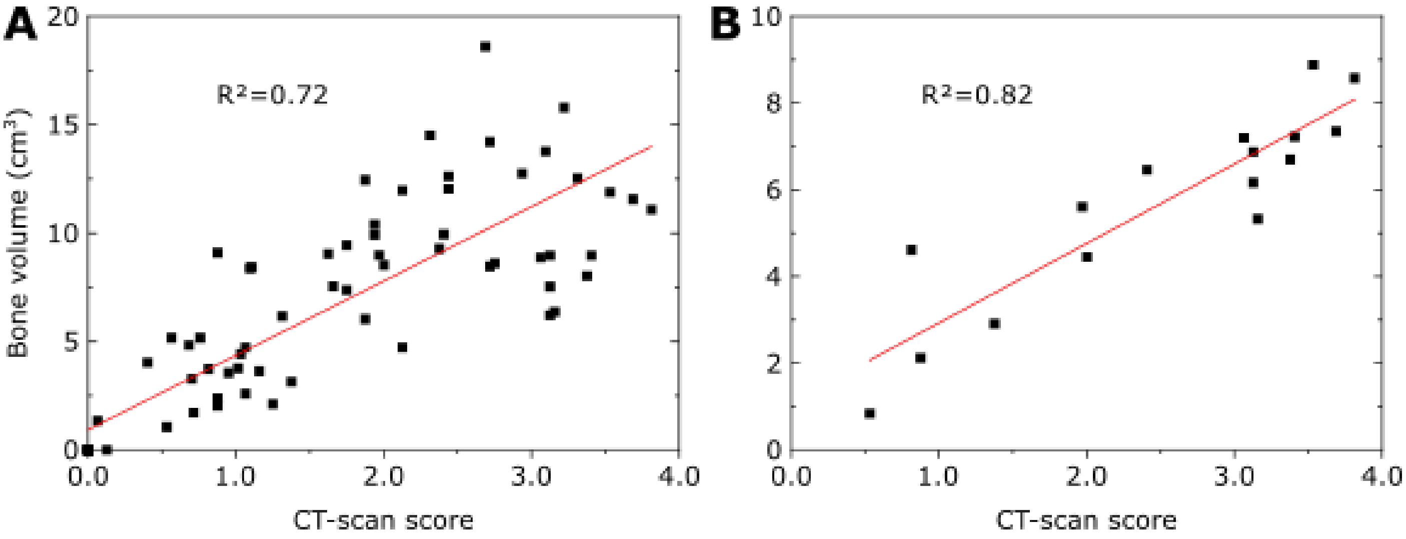


**Figure SI 10.** Correlation between the CT-scan score given by the clinicians and the quantitative analyses of new bone volume expressed in $cm^3$. (A) Correlation between the new bone volume measured using the CT scans and the CT-scan score. A linear tendency is to be observed and the correlation coefficient $R^2$ is given. (B) Correlation between the quantification of the new bone volume using μCT acquisitions and the CT-scan score. A linear fit was drawn with a good correlation coefficient $R^2$.
